# ParticleChromo3D: A Particle Swarm Optimization Algorithm for Chromosome and Genome 3D Structure Prediction from Hi-C Data

**DOI:** 10.1101/2021.02.11.430871

**Authors:** David Vadnais, Michael Middleton, Oluwatosin Oluwadare

## Abstract

The three-dimensional (3D) structure of chromatin has a massive effect on its function. Because of this, it is desirable to have an understanding of the 3D structural organization of chromatin. To gain greater insight into the spatial organization of chromosomes and genomes and the functions they perform, chromosome conformation capture techniques, particularly Hi-C, have been developed. The Hi-C technology is widely used and well-known because of its ability to profile interactions for all read pairs in an entire genome. The advent of Hi-C has greatly expanded our understanding of the 3D genome, genome folding, gene regulation and has enabled the development of many 3D chromosome structure reconstruction methods. Here, we propose a novel approach for 3D chromosome and genome structure reconstruction from Hi-C data using Particle Swarm Optimization approach called ParticleChromo3D. This algorithm begins with a grouping of candidate solution locations for each chromosome bin, according to the particle swarm algorithm, and then iterates its position towards a global best candidate solution. While moving towards the optimal global solution, each candidate solution or particle uses its own local best information and a randomizer to choose its path. Using several metrics to validate our results, we show that ParticleChromo3D produces a robust and rigorous representation of the 3D structure for input Hi-C data. We evaluated our algorithm on simulated and real Hi-C data in this work. Our results show that ParticleChromo3D is more accurate than most of the existing algorithms for 3D structure reconstruction. Our results also show that constructed ParticleChromo3D structures are very consistent, hence indicating that it will always arrive at the global solution at every iteration. The source code for ParticleChromo3D, the simulated and real Hi-C datasets, and the models generated for these datasets are available here: https://github.com/OluwadareLab/ParticleChromo3D

## Introduction

Chromosome Conformation Capture (3C) and its subsequent derivative technologies are invaluable for describing chromatin’s three-dimensional (3D) structure [1]. 3C’s biochemical approach to studying DNA’s topography within chromatin has outperformed the traditional microscopy approaches like fluorescence in situ hybridization (FISH) due to 3C’s systematic nature [2]. As a side note, Microscopy is still used in conjunction with 3C for verifying the actual 3D structure of chromatin against the predicted outcome [1]. 3C was first described by [3] Dekker et al. (2002). Since then, more technologies were developed [4], such as the Chromosome Conformation Capture-on-Chip (4C) [5], Chromosome Conformation Capture Carbon Copy (5C) [6], Hi-C[7], TCC[8], and Chromatin Interaction Analysis by Paired-End Tag sequencing ChIA-PET [2,9]. These derivative technologies were designed to augment 3C’s in the following areas, measure spatial data within chromatin, increase measuring throughput, and analyze proteins and RNA within chromatin instead of just DNA. Lieberman-Aiden et al., 2009 [7] designed Hi-C as a minimally biased “all vs. all” approach. Hi-C works by injecting biotin-labeled nucleotides during the ligation step [4]. Hi-C provides a method for finding genome-wide chromatin IF data in the form of a contact matrix [1].

Hi-C analysis doubtlessly introduced great benefit to 3D genome research— they explain a series of events such as genome folding, gene regulation, genome stability, and the relationship between regulatory elements and structural features in the cell nucleus [2,7,10]. Importantly, it is possible to glean insight into chromatin’s 3D structure using the Hi-C data. However, to use Hi-C data for 3D structure modeling, some pre-processing is necessary to extract the interaction frequencies (IF) between the chromosome or genome’s interacting loci [11]. This process involves quality control and mapping of the data [12]. Once these steps are completed, an IF matrix, or called contact matrix or map, is generated. An IF matrix is a symmetric matrix that records a one-to-one interaction frequency for all the intersecting loci [7,10]. The IF matrix is represented as either a square contact matrix or as a three-column sparse matrix. Each cell has genomic bins within these matrices that are the length of the data’s resolution representing each cell [12]. Hence, the higher the resolution (5KB), the larger the contact matrix’s size. And similarly, the lower the resolution (1MB), the smaller the contact matrix’s size. Next, this Hi-C data is normalized to remove biases that next-generation sequencing can create [12,13]. An example of this type of bias would be copy number variation [13]. Other systematic biases introduced during the Hi-C experiment are by external factors, such as DNA shearing and cutting [10]. Today, several computational algorithms have been developed to remove these biases from the Hi-C IF data [13–20]. Once the Hi-C IF matrix data is normalized, it is most suitable for 3D chromosome or genome modeling. Some tools have been developed to automate this Hi-C pre-processing steps; they include GenomeFlow [21], Hi-Cpipe [22], Juicer [23], HiC-Pro [24], and HiCUP [25].

To create 3D chromosome and genome structures from IF data, many techniques can be used. Oluwadare, O., *et al.* (2019) [10] pooled the various developed analysis techniques into three buckets, which are Distance-based, Contact-based, and Probability-based methods. The first method is a Distance-based method that maps IF data to distance data and then uses an optimizer to solve for the 3D coordinates [12]. This type of analysis’s final output will be (x, y, z) coordinates [12]. However, the difficulty is picking out how to convert the IF data and which optimization algorithm to use [10]. The distance between two genomic bins is often represented as *Distance_i,j_*= 1/(*IF*_*i*,*j*_^*α*^) [10,11]. In this approach *IF*_*i*,*j*_ is the number of times two genomic bins had contact and *α* is a factor which is used for modeling, called the conversion factor. This distance can then be optimized against other genomic bins’ other distance values to create a 3D model. Several methods [10] belong in this category include, ChromSDE[26], AutoChrom3D [27], Chromosome3D [28], 3DMax [29], ShRec3D [30], LorDG [31], InfMod3DGen [32], HSA [33], ShNeigh[34]. The second classification for 3D genome structure modeling algorithms from IF data is Contact-based methods. This technique uses the IF data directly instead of starting by converting the data to a (x, y, z) coordinate system [10]. One way to model this data is with a gradient descent/ascent algorithm [10]. This approach was explored by Trieu T, and Cheng J., 2015 through the algorithm titled MOGEN [35]. MOGEN works by optimizing a scoring function that scores how well the chromosomal contact rules have been satisfied [35]. Another contact method was to take the interaction frequency and use it for spatial restraints [36]. Gen3D [37], Chrom3D [38], and GEM [39] are other examples in this category. The third classification is Probability-based. The advantages of probability-based approaches are that they easily account for uncertainties in experimental data and can perform statistical calculations of noise sources or specific structural properties [10]. Unfortunately, probability techniques can be very time-consuming compared to Contact and Distance methods. Rousseau et al., 2011 created the first model in this category using a Markov chain Monte Carlo approach called MCMC5C [40]. Markov chain Monte Carlo was used due to its synergy with estimating properties’ distribution [10]. Varoquaux. N., et al., 2014 [41] extended this probability-based approach to modeling the 3D structure of DNA. They used a Poisson model and maximized a log-likelihood function [41]. Many other statistical models can still be explored.

This paper presents ParticleChromo3D, a new distance-based algorithm for chromosome 3D structure reconstruction from Hi-C data. ParticleChromo3D uses Particle Swarm Optimization (PSO) to generate 3D structures of chromosomes from Hi-C data. Here, we show that ParticleChromo3D can generate candidate structures for chromosomes from Hi-C data. Additionally, we analyze the effects of parameters such as confidence coefficient and swarm size on the structural accuracy of our algorithm. Finally, we compared ParticleChromo3D to a set of commonly used chromosome 3D reconstruction methods, and it performed better than most of these methods. We showed that ParticleChromo3D effectively generates 3Dstructures from Hi-C data and is highly consistent in its modeling performance.

## Materials and Methods

### The Particle Swarm Optimization Algorithm

Kennedy J., and Eberhart R. (1995) [42] developed the Particle Swarm Optimization (PSO) as an algorithm that attempts to solve optimization problems by mimicking the behavior of a flock of birds. PSO has been used in the following fields: antennas, biomedical, city design/civil engineering, communication networks, combinatorial optimization, control, intrusion detection/cybersecurity, distribution networks, electronics and electromagnetics, engines and motors, entertainment, diagnosis of faults, the financial industry, fuzzy logic, computer graphics/visualization, metallurgy, neural networks, prediction and forecasting, power plants, robotics, scheduling, security and military, sensor networks, and signal processing [43–47]. Since PSO has been used in so many disparate fields, it appears to be robust and flexible, which gives credence to the idea that it could be used in this use case of bioinformatics and many others [48]. PSO falls into the optimization taxonomy of swarm intelligence [49]. PSO works by creating a set of particles or actors that explore a topology and look for the global minimum of that topology [49]. At each iteration, the swarm stores each particle’s minimum result, as well as the global swarm’s minimum, found. The particles explore the space with both a position and velocity, and they change their velocity based on three parameters. These three parameters are current velocity, distance to the personal best, and distance to the global best [49]. Position changes are made based on the calculated velocity during each iteration. The velocity function is as follows [50]:

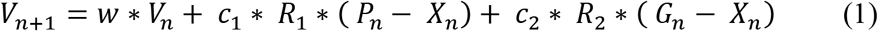

Then position is updated as follow:

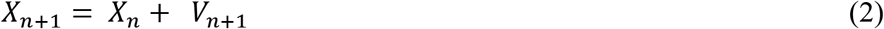

Where:

- *V*_*n*_ is the current velocity at iteration *n*
- *c*_1_ and *c*_2_ are two real numbers that stand for local and global weights and are the personal best of the specific particle and the global best vectors, respectively, at iteration *n* [50].
- The *R*_1_ and *R*_2_ values are randomized values used to increase the explored terrain [50].
- *w* is the inertia weight parameter, and it determines the rate of contribution of a velocity [42].
- *G*_*n*_ represents the best position of the swarm at iteration *n*.
- *P*_*n*_ represents the best position of an individual particle.
- *X*_*n*_ is the best position of an individual particle at the iteration *n*.

### Why PSO

This project’s rationale is that using PSO could be a very efficient method for optimizing Hi-C data due to its inherent ability to hold local minima within its particles. This inherent property will allow sub-structures to be analyzed for optimality independently of the entire structure.

In Fig 1, particle one is at the global best minimum found so far. However, particle two has a better structure in its top half, and it is potentially independent of the bottom half. Because particle one has a better solution so far, particle two will traverse towards the structure in particle one in the iteration *n* + 1. While particle two is traversing, it will go along a path that maintains its superior 3D model sections. Thus, it has a higher chance of finding the absolute minimum distance value. The more particles there are, the greater the time complexity of PSO and the higher the chance of finding the absolute minimum. The inherent breaking up of the problem could lend itself to powerful 3D structure creation results. More abstractly relative to Hi-C data but in the traditional PSO sense, the same problem as above might look as follows (Fig 2) when presented in a topological map.

**Fig 1.**
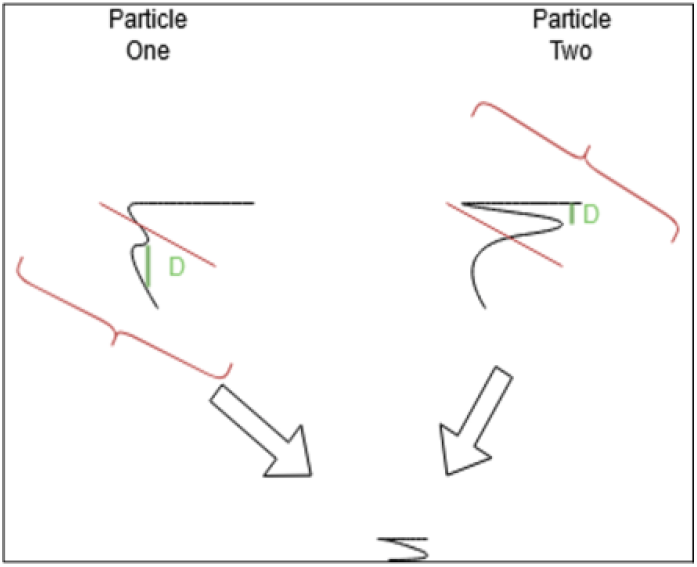
PSO potential advantage for structure holding. The figure summarizes the PSO algorithm performance expectation on the 3D genome structure reconstruction problem.

**Fig 2.**
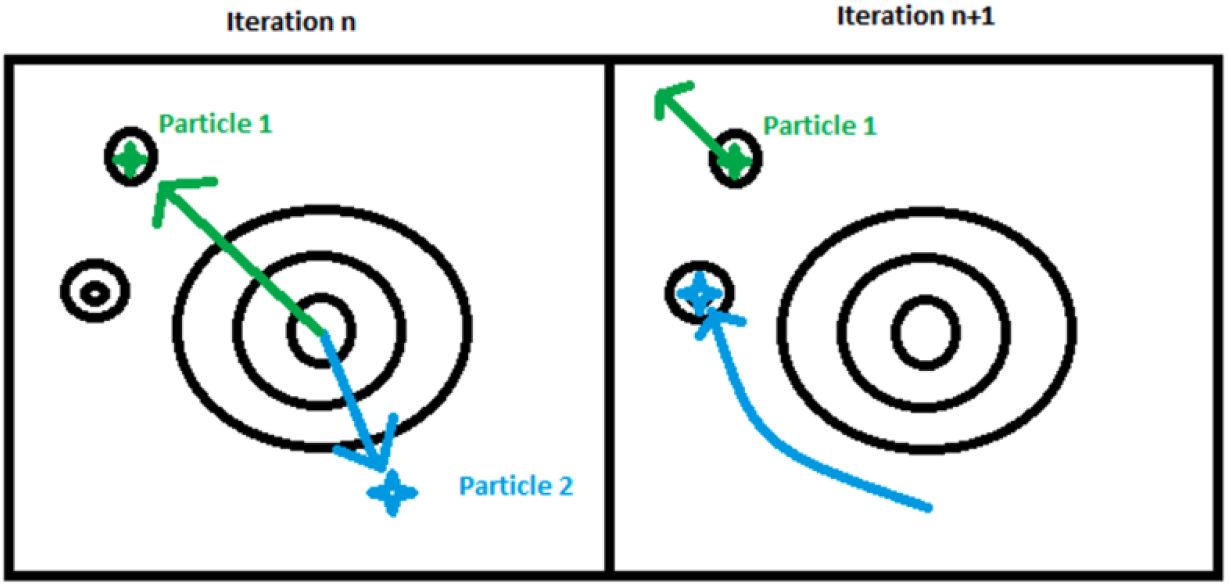
PSO particle iteration description. This figure explains the PSO algorithm’s search mechanism for determining the best 3D structure following the individual particles’ modified velocity and position in the swarm.

From Fig 2, in the *nth* iteration, particle 1 found a local minimum within this step. Since of all the particles, this is the lowest point; particle two will search towards particle one with a random chance amount added to its velocity [49, 51, 52]. The random chance keeps particle two from going straight to the optimal solution [49,53]]. In this case, particle two found the absolute minima, and from here on, all the particles will begin to migrate towards particle 2. We will test this hypothesis by analyzing its output with the evaluation metrics defined in the “Results” section.

In summary, we believe the particle-based structure of PSO may lend itself well to the problem of converting Hi-C IF data into 3D models. We will test this hypothesis and compare our results to the existing modeling methods.

### PSO for 3D Structure Reconstruction from Hi-C Data

Here we describe how we implemented the PSO algorithm as a distance-based approach for 3D genome reconstruction from Hi-C data. This algorithm is called ParticleChromo3D. In this context, the input IF data is converted to the distance equivalent using the conversion factor, *α*, for 3D structure reconstruction. First, we initialize the particles’ 3D (x,y,z) coordinates for each genomic bin or regions randomly in the range [-1, 1]. We used the sum of squared error function as the loss function to compute chromosome structures from a contact map. Finally, we used PSO to iteratively improve our function until it has converged on either an absolute or local minima. The full ParticleChromo3D algorithm is presented in Fig 3. Some parameters are needed to use the PSO algorithm for 3D structure reconstruction. This work has provided the parameter values that produced our algorithm’s optimal results. The users can also provide their settings to fit their data where necessary. The results of the series of tests and validation performed to determine the default parameters are described in the “Parameters Estimation” section of the Results section.

**Fig 3.**
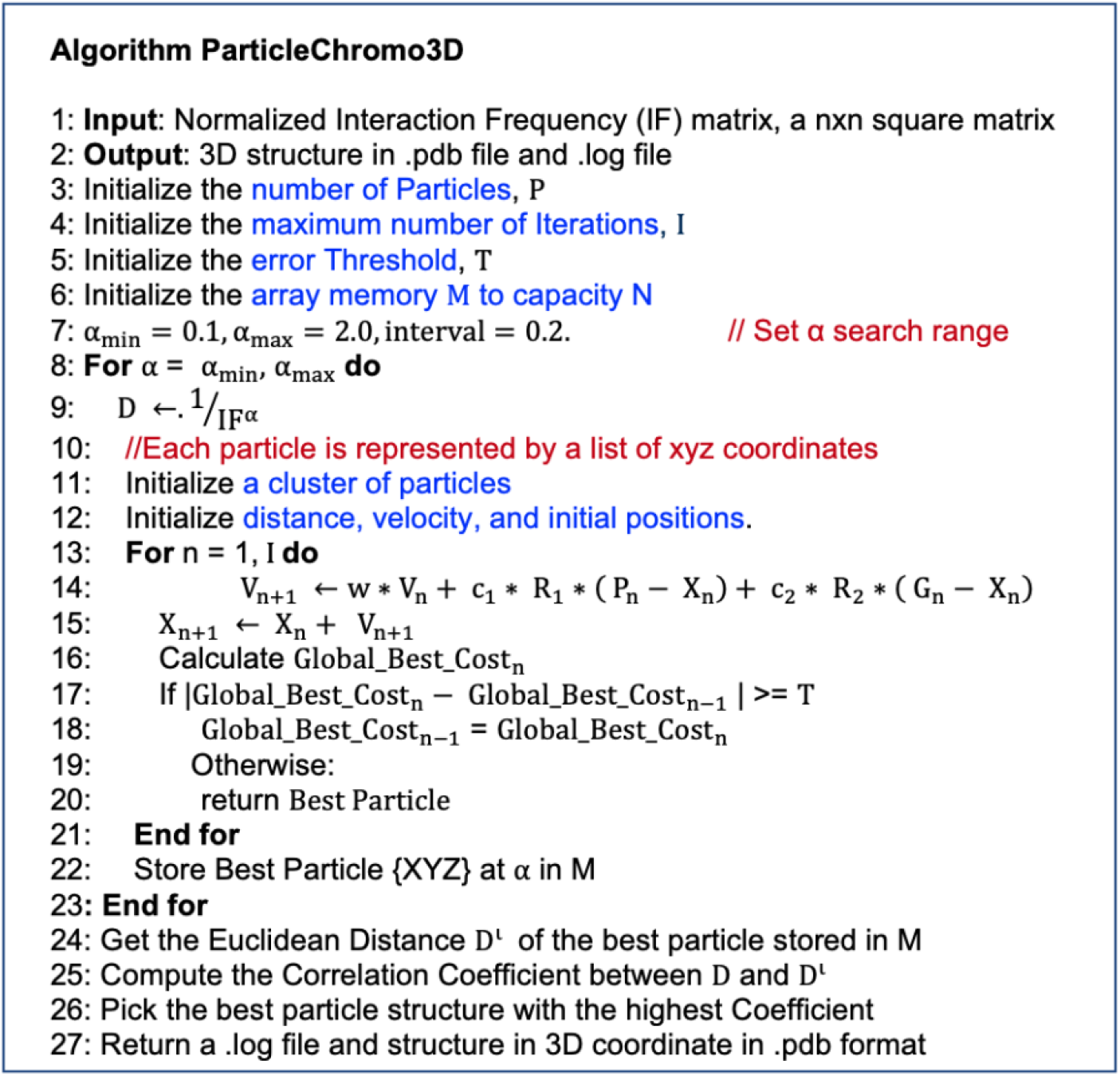
PSO for chromosome and genome 3D Structure prediction. We present a step-by-step illustration of the significant steps taken by ParticleChromo3D for 3D chromosome and genome structure reconstruction from an input normalized IF matrix.

### Model Representation

A particle is a candidate solution. A list of XYZ coordinates represents each particle in the solution. The candidate solution’s length in the number of regions in the input Hi-C data. Each particle’s point is the individual coordinate, XYZ, of each bead. A swarm consists of N candidate solution, also called the swarm size, which the user provides as program input. We provide more explanation in the “Parameters Estimation” section below for how to determine the swarm size.

### Data

Our study used the yeast synthetic or simulated dataset from Adhikari et al., 2016 [28] to perform parameter tuning and validation. The simulated dataset was created from a yeast structure for chromosome 4 at 50kb resolution [54]. The number of genome loci in the synthetic dataset is 610. We used the GM12878 cell Hi-C dataset to analyze a real dataset, GEO Accession number GSE63525[55]. The normalized contact matrix was downloaded from the GSDB database with GSDB ID: OO7429SF [56].

## Results

### Metrics Used for Evaluation

To evaluate the structure’s consistency with the input Hi-C matrix, we used the following metrics:

### Pearson Correlation Coefficient (PCC)

The Pearson correlation coefficient is as follows [10],

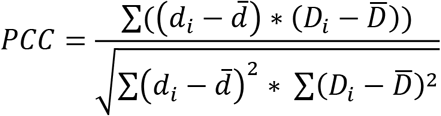

Where:

- *D*_*i*_ and *d*_*i*_ are instances of a distance value between two bins.
- 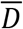 and *d̅* are the means of the distances within the data set.
- It measures the relationship between variables. Values a between −1 to +1
- A higher value is better.

### Spearman Correlation Coefficient (SCC)

Spearman’s correlation coefficient is defined below [10],

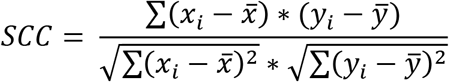

Where:

- x_i_ and y_i_ are the rank of the distances,*D*_*i*_ and *d*_*i*_, defined in the PCC equation above.
- 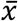 and 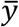 are the sample mean tank of both x and y, respectively.
- Values a between −1 to +1. A higher value is better.

### Root Mean Squared Error (RMSE)

Root mean squared error follows the equation below [10],

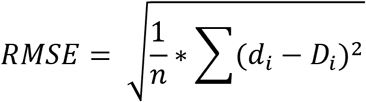

Where:

- Di and di are instances of distance values from the data and another data source.
- The value n is the size of the data set.

### TM-Score

TM-Score is defined as follows [42][43],

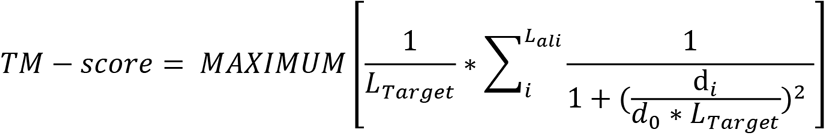

Where:

- L_Target_ is the length of the chromosome.
- di is an instance of a distance value between two bins.
- L_ali_ Represents the count of all aligned residues.
- d_0_ is a normalizing parameter.

The TM-score is a metric to measure the structural similarity of two proteins or models [57,58]. A TM-score value can be between (0,1] were 1 indicates two identical structures [57]. A score of 0.17 indicates pure randomness, and a score above 0.5 indicates the two structures have mostly the same folds [58]. Hence the higher, the better.

### Parameters Estimation

We used the yeast synthetic dataset to decide on ParticleChromo3D’s best parameters. We used this data set to investigate the mechanism for choosing the best alpha conversion factor for input Hi-C data. Also, determine the optimal swarm size; determine the best threshold value for the algorithm, inertia value(w), and the best coefficients for our PSO velocity (*c*_1_ and *c*_2_). We evaluated our reconstructed structures by comparing them with the synthetic dataset’s true distance structure provided by Adhikari et al., 2016 [28]. We evaluated our algorithms with the PCC, SCC, RMSE, and TM-score metrics. Based on the results from the evaluation, the default value for the ParticleChromo3D parameters are set as presented below:

### Conversion Factor Test (*α*)

The synthetic interaction frequency data set was generated from a yeast structure for chromosome 4 at 50kb [69] with an *α* value of 1 using the formula: *IF* = 1/*D*^*α*^. Hence, the relevance of using this test data is to test if our algorithm can predict the alpha value used to produce the synthetic dataset. For both PCC and SCC, our algorithm performed best at a conversion factor (alpha) of 1.0 (Fig 4). Our algorithm’s default parameter setting is that it searches for the best alpha value in the range [0.1, 1.5]. Side by side comparison of the true simulated data (yeast) structure and the reconstructed structure by ParticleChromo3D shows that they are highly similar (Fig 5)

**Fig 4.**
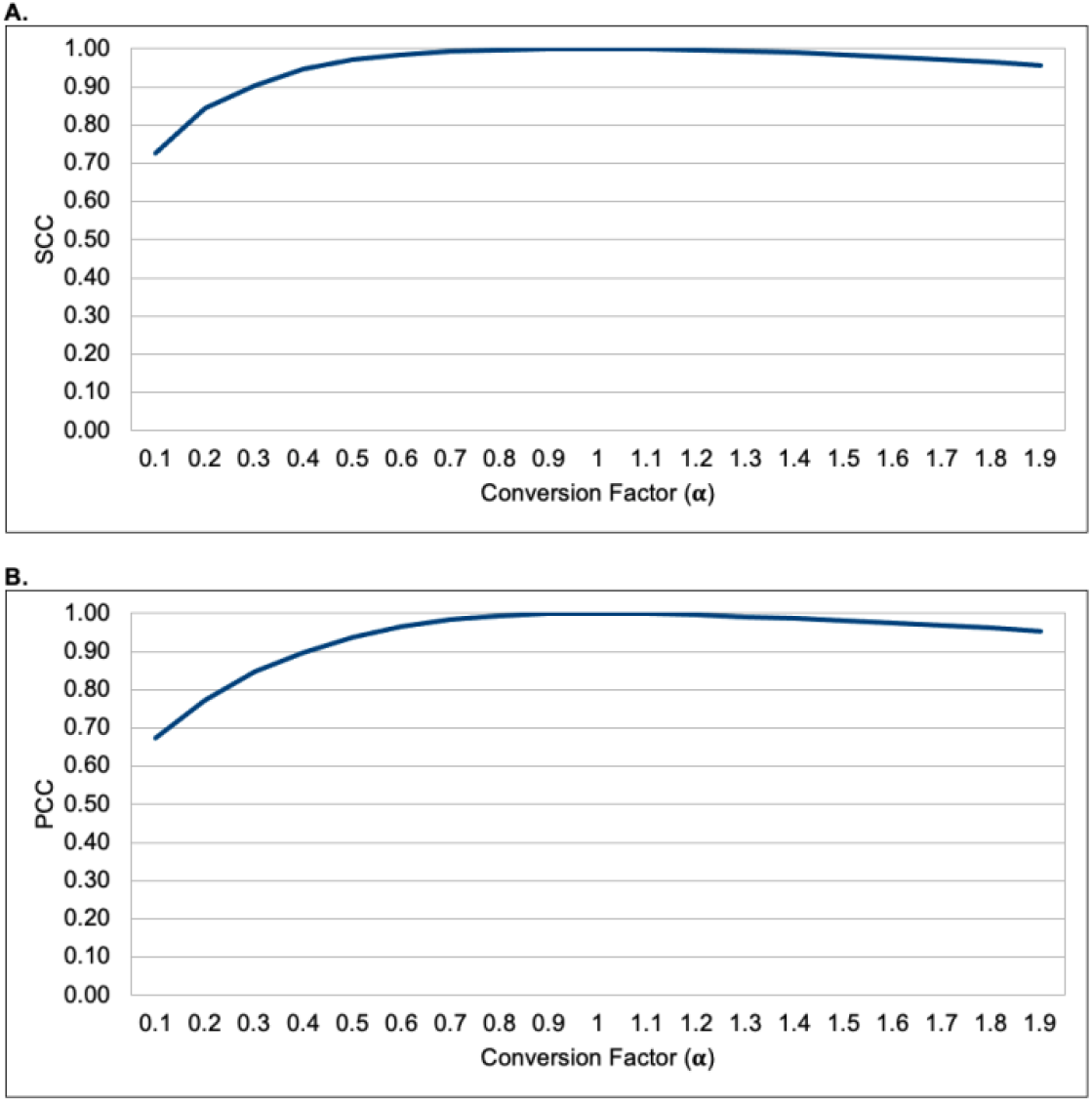
A plot of the evaluation metric versus the conversion factors. (A) A plot of SCC vs. Conversion factor. (B) A plot of PCC vs. Conversion factor. Here, we show the performance of ParticleChromo3D on the SCC and PCC metric for the simulated dataset at *α* value in the range 0.1 to 1.9. The result shows the best result is recorded at *α* = 1. The SCC and PCC metric values were obtained by comparing the ParticleChromo3D algorithm’s output structure at each *α* value with the true structure. In Fig 4A and 4B, the Y-axis denotes the SCC and PCC scores, respectively, in the range [-1,1], and the X-axis denotes the conversion factor values. A higher SCC and PCC value is better.

**Fig 5.**
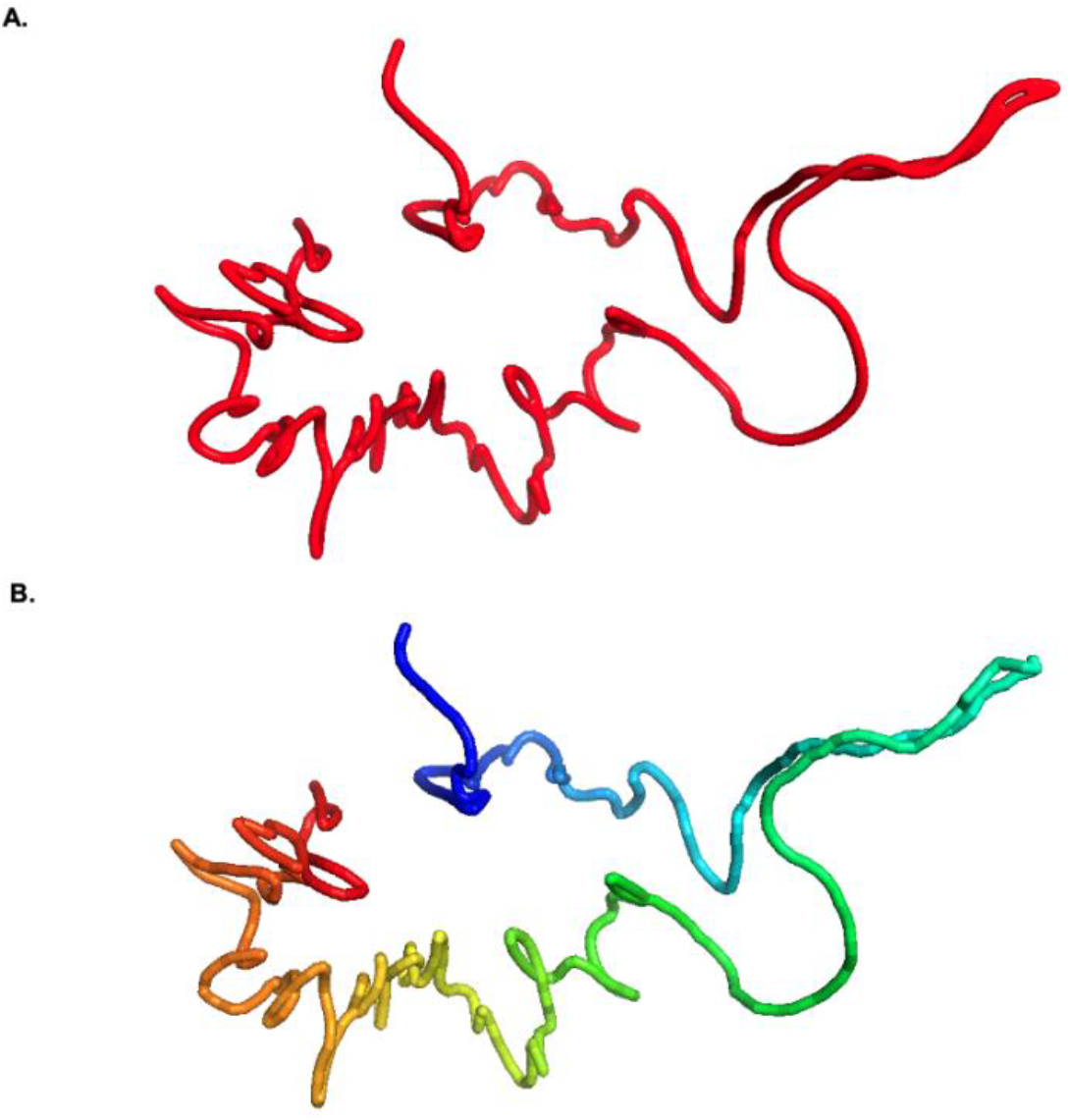
A comparison of the simulated data true structure and reconstructed structure by ParticleChromo3D. (A)True structure from Duan et al. [54] (B) Reconstructed structures for the simulated data using ParticleChromo3D.

### Swarm Size

The swarm size defines the number of particles in the PSO algorithm. We evaluated the performance of the ParticleChromo3D with changes in swarm size (Fig 6A, Fig 6B, Fig 6C). Also, we evaluated the effect of an increase in swarm size against computation time (Fig 6D). Our result shows that computational time increases with increased swarm size. Given the computational implication and the algorithm’s performance at various swarm size, we defined a swarm size of 15 as our default value for this parameter. According to our experiments, the Swarm size 10 is most suitable if the user’s priority is saving computational time, and swarm size 20 is suitable when the user’s preference is algorithm performance over time. Hence, setting the default swarm size 15 gives us the best of both worlds. The structures generated by ParticleChromo3D also shows that the result at swarm size 15(Fig 7C) and 20(Fig 7D) are most similar to the simulated data true structure represented in Fig 5A (Fig 7).

**Fig 6.**
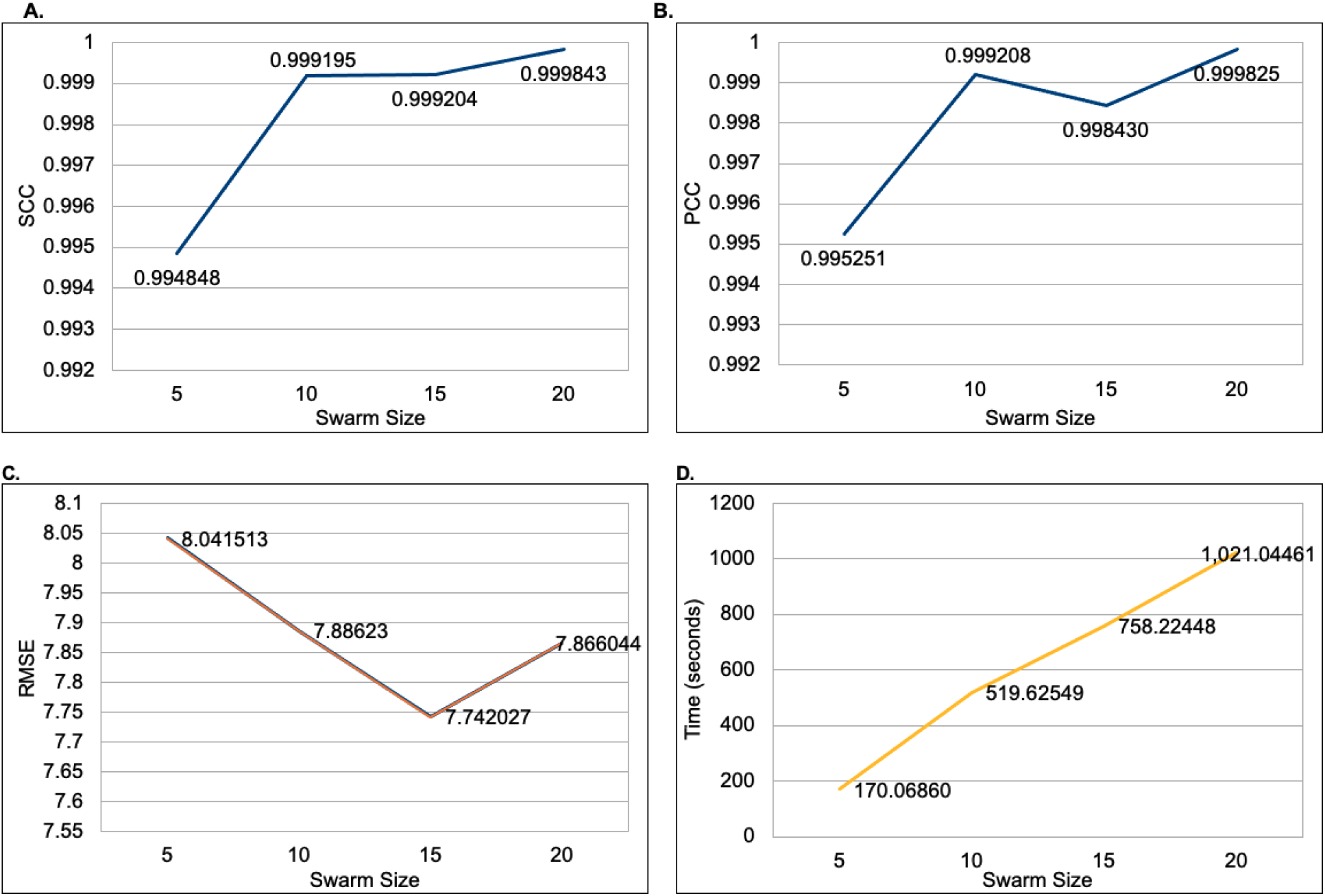
A plot of the evaluation metric versus the Swarm Size parameter. (A) A plot of the SCC vs. the Swarm Size. (B) A plot of PCC vs. the Swarm Size. (C) A plot of RMSE vs. the Swarm Size. (D) A plot of the runtime, in seconds, vs. the Swarm Size. The SCC, PCC, and RMSE values were obtained by comparing the ParticleChromo3D algorithm’s output structure with the simulated data true structure. In Fig 6A and Fig 6B, the Y-axis denotes the SCC and PCC score in the range [-1,1], and the X-axis denotes the Swarm Sizes values considered. A higher SCC and PCC value is better. In Fig 6C, the Y-axis denotes the RMSE score, and the X-axis denotes the Swarm Size values. A lower RMSE value is better. In Fig 6D, the Y-axis denotes the running time in seconds, and the X-axis denotes the Swarm Size values.

**Fig 7.**
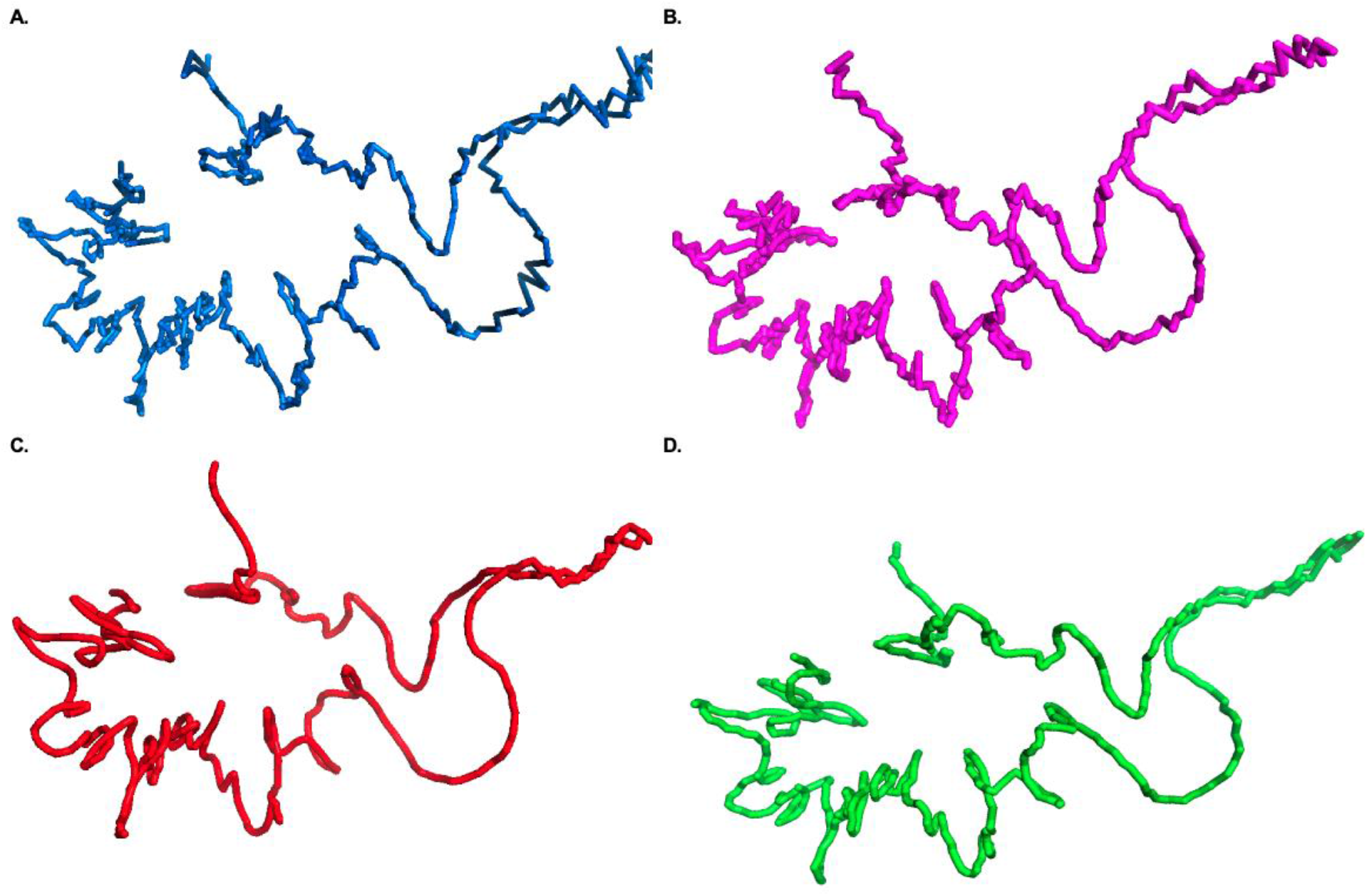
Structures generated by ParticleChromo3D at different swarm size values. Here, we show the structure generated at Swarm size = (A) 5 (Represented with blue color), (B) 10 (Represented with magenta color), (C) 15 (Represented with red color), and (D) 20 (Represented with green color). As shown, the structure generated at swarm size 5 is not smooth; it has a couple of rough edges (Fig 7A). This correlates to the SCC, PCC, and RMSE recorded at this swarm size as it is the lowest at swarm size 5. Next, at Swarm size 10(Fig 7B), we observe a smoother representation but still with some rough edges. The result here shows that the results were really similar at swarm size 15 and 20(Fig 7C, Fig 7D).

### Threshold

The threshold parameter is designed to serve as an early stopping criterion if the algorithm converges before the maximum number of iterations is reached. Hence, we evaluated the effect of varying threshold levels using the evaluation metrics(Fig 8). The output structures generated by each of the thresholds also allow a visual examination of a threshold value(Fig 9). We observed that the lower the threshold, the more accurate(Fig 8) and similar the structure is to the generated true simulated data structure in Fig 5A(Fig 9F). It worth noting that this does have a running time implication. Reducing the threshold led to a longer running time. However, since this was a trade-off between a superior result and longer running time or a fairly good result and short running time, we chose the former for ParticleChromo3D. The default threshold for our algorithm is 0.000001.

**Fig 8.**
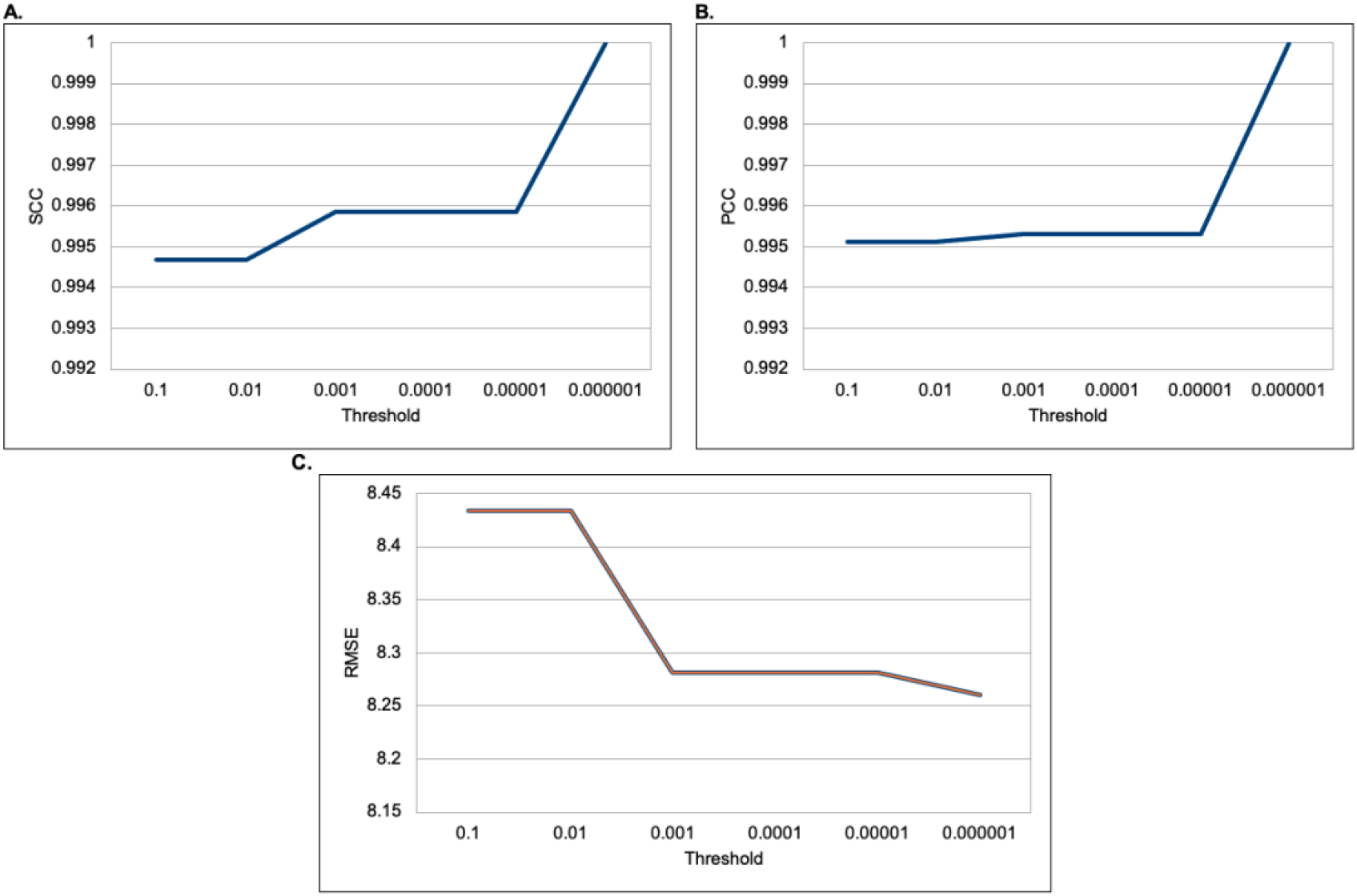
A plot of the evaluation metric versus the Threshold parameter. (A) A plot of the SCC versus different threshold levels. (B) A plot of PCC versus the different threshold levels. (C) A plot of RMSE versus different threshold levels. The results show the performance of our algorithm at threshold values 0.1, 0.01, 0.001, 0.0001, 0.00001, 0.000001. The SCC, PCC, and RMSE values reported were obtained by comparing the ParticleChromo3D algorithm’s output structure to the simulated dataset’s true structure. In Fig 8A and 8B, the Y-axis denotes the SCC and PCC scores in the range [-1,1], and the X-axis denotes the Threshold values. A higher SCC and PCC value is better. In Fig 8C, Y-axis denotes the RMSE score, and the X-axis denotes the Threshold values. A lower RMSE value is better.

**Fig 9.**
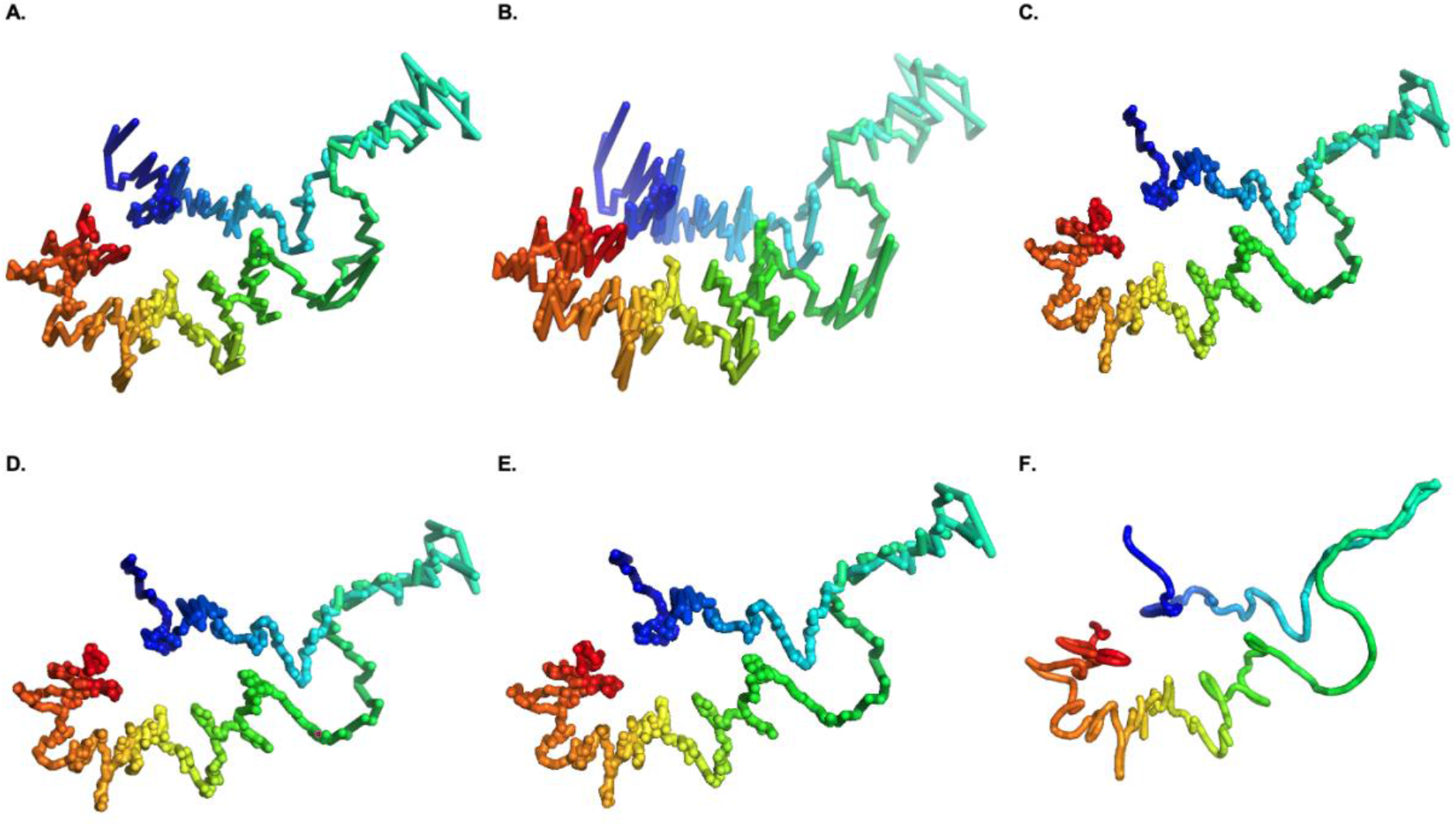
Structures at a threshold of 0.1, 0.01, 0.001, 0.0001, 0.00001, 0.000001 respectively. (A) Represents the structure produced using a threshold of 0.1. (B) Represents the structure produced using a threshold of 0.01. (C) Represents the structure produced using a threshold of 0.001. (D) Represents the structure produced using a threshold of 0.0001. (E) Represents the structure produced using a threshold of 0.00001. (F) Represents the structure produced using a threshold of 0.000001. The results showed that the threshold value of 0.000001, Fig 9F, produced the best result.

### Confidence Coefficient (*c*_1_ and *c*_2_)

The *c*_1_ and *c*_2_ parameters represent the local-confidence and local and global swarm confidence level coefficient. Kennedy and Eberhart, 1995[42] proposed that *c*_1_**=***c*_2_ = 2. We experimented with testing how this value’s changes affected our algorithm’s accuracy for local confidence coefficient (*c*_1_) 0.3 to 0.9 and global confidence values 0.1 to 2.8 (S1 and S2 Fig). From our results, we found that a local confidence coefficient (*c*_1_) of 0.3 with a global confidence coefficient (*c*_2_) of 2.5 performed best (Fig 10). Hence, these values were set as ParticleChromo3D’s confidence coefficient values. The accuracy results generated for all the local confidence coefficient (c1) at varying global confidence values is compiled in Fig 11.

**Fig 10.**
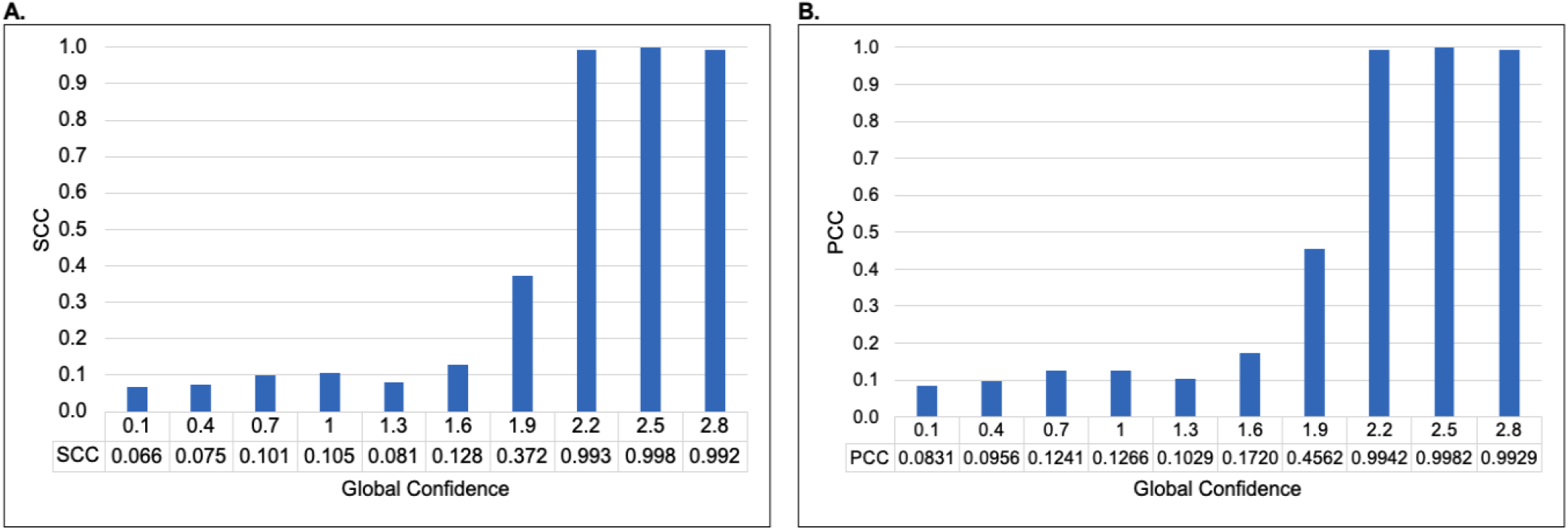
Confidence Coefficient Test. (A) A plot of the SCC by Global Confidence at Local Confidence 0.3. (B) A plot of PCC by Global Confidence at Local Confidence 0.3. The plot of the local confidence value local confidence coefficient (*c*_1_) = 0.3 against the varying level of global confidence coefficient (*c*_2_) values from 0.1 to 2.8. The results show that the best result was obtained at *c*_2_ = 2.5. The SCC and PCC values reported were obtained by comparing the ParticleChromo3D algorithm’s output structure with the simulated dataset’s true structure. In Fig 10A and 10B, the Y-axis denotes the SCC and PCC scores in the range [-1,1], the X-axis denotes the global confidence values, and the colored plot denotes the local confidence values. A higher SCC and PCC value is better.

**Fig 11.**
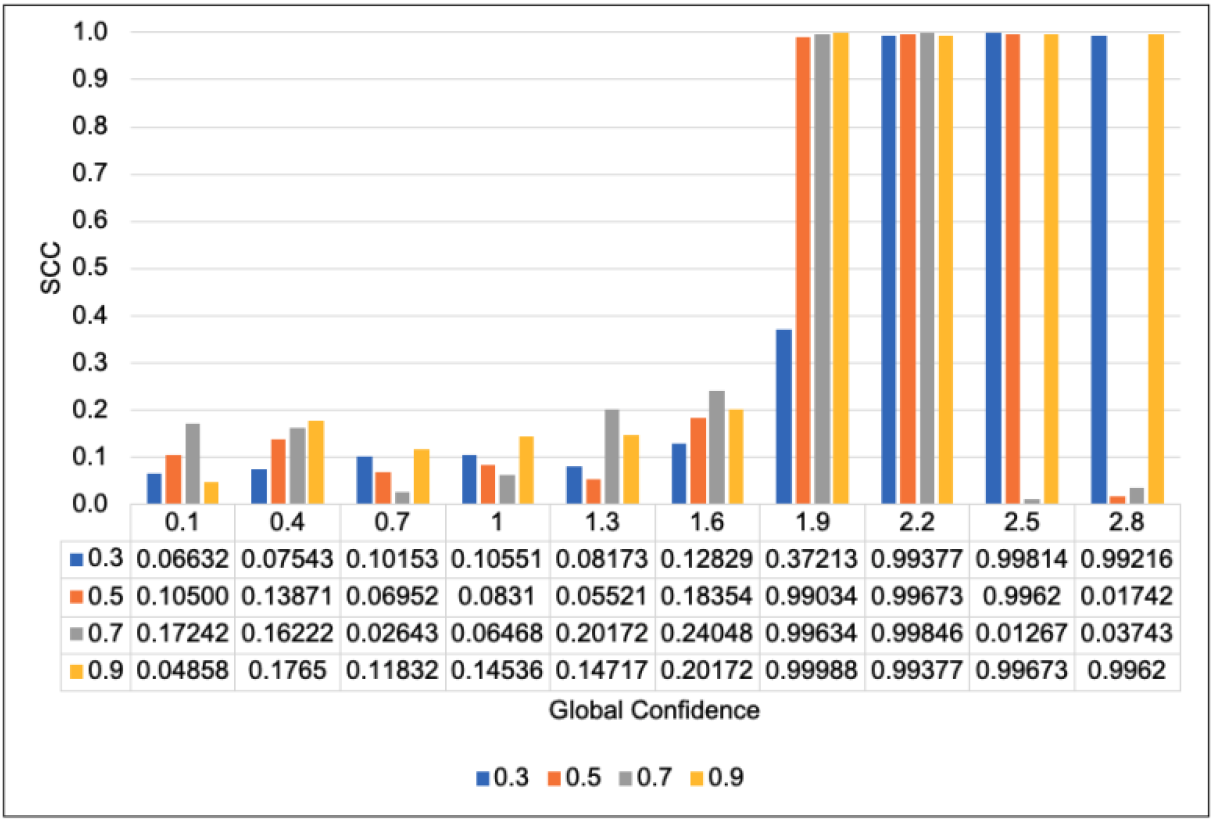
A combined plot of different local confidences versus global confidences. The result’s combined plot was obtained by comparing the ParticleChromo3D algorithm’s output structure with the simulated dataset’s true structure for local confidence values of 0.3 to 0.9 and global confidence values of 0.1 to 2.8. This plot shows the SCC accuracy of the structures generated. The Y-axis denotes the SCC score in the range [-1,1], the X-axis denotes the global confidence values, and the colored plot denotes the local confidence values. A higher SCC value is better.

### Random Numbers (*R*_1_ and *R*_2_)

*R*_1_ and *R*_2_ are uniform random numbers between 0 and 1[59].

### Assessment on simulated data

We evaluated how noise levels affect ParticleChromo3D’s ability to predict chromosome 3D structures in the presence of noise. Using the yeast synthetic dataset from Adhikari et al., 2016 [28]. The data were simulated with a varying noise level. Adhikari, et al. introduced noise into the yeast IF matrix to make 12 additional datasets with different levels of noise at 3%, 5%, 7%, 10%, 13%, 15%, 17%, 20%, 25%, 30%, 35%, and 40%. As reported by the authors, converting this IF to their distance equivalent produced distorted distances that didn’t match the true distances. They were thereby simulating the inconsistent constraints that can sometimes be observed in un-normalized Hi-C data. As shown, our algorithm performed the best with no noise in the data at 0 (Fig 12).

**Fig 12.**
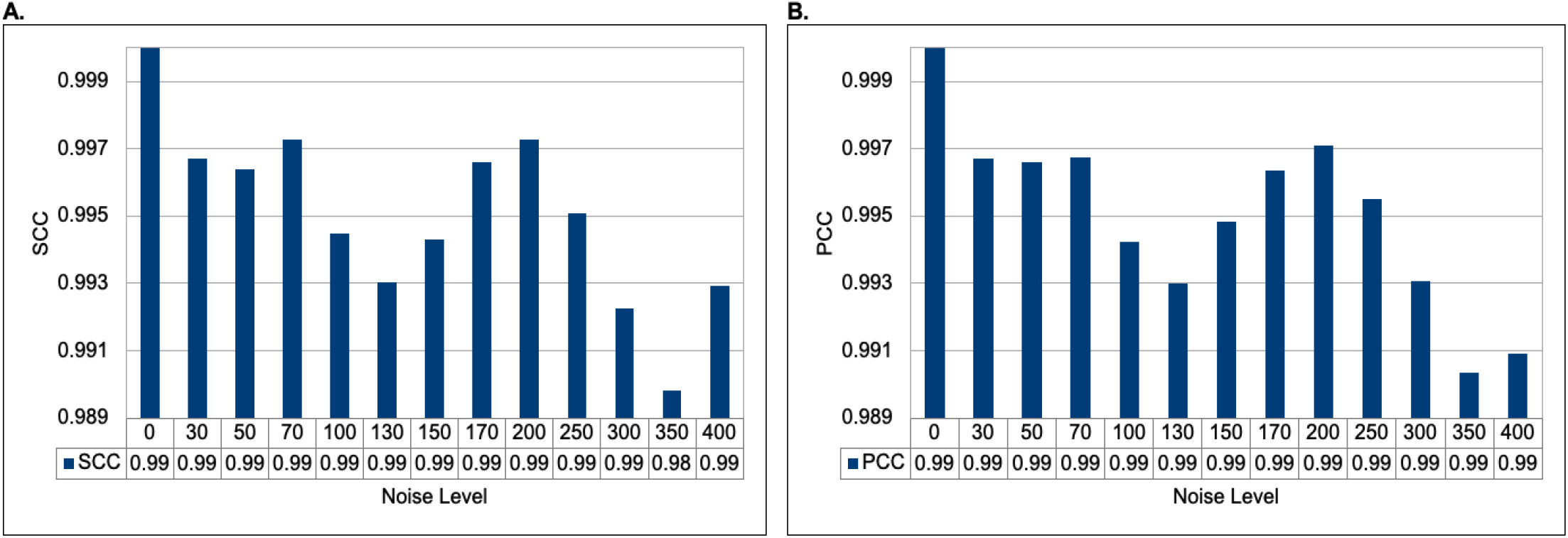
Assessment of the structures generated by ParticleChromo3D for the simulated dataset on varying noise levels. (A) A plot of the SCC versus Noise level. (B) A plot of PCC versus the Noise level. This plot shows the SCC and PCC accuracy of the structures generated by ParticleChromo3D at different noise levels introduced. In Fig 12A and Fig 12B, the Y-axis denotes the metric score in the range [-1,1]. The X-axis denotes the Noise level. A higher SCC and PCC value is better.

Furthermore, the other result obtained by comparing the ParticleChromo3D algorithm’s output structure from the noisy input datasets with the simulated dataset’s true structure shows that it can achieve a competitive result when dealing with un-normalized or noisy Hi-C datasets(Fig 13). The result shows that our algorithm can achieve the results obtainable at reduced noise level even at increased noise as indicated by Noise 7%(Fig 13B) and 20%(Fig 13C), respectively (Fig 12). Also, the difference in performance between the best structure and the worst structure is ~0.01. Hence, our algorithm cannot be potentially be affected by the presence of noise in the input Hi-C data.

**Fig 13.**
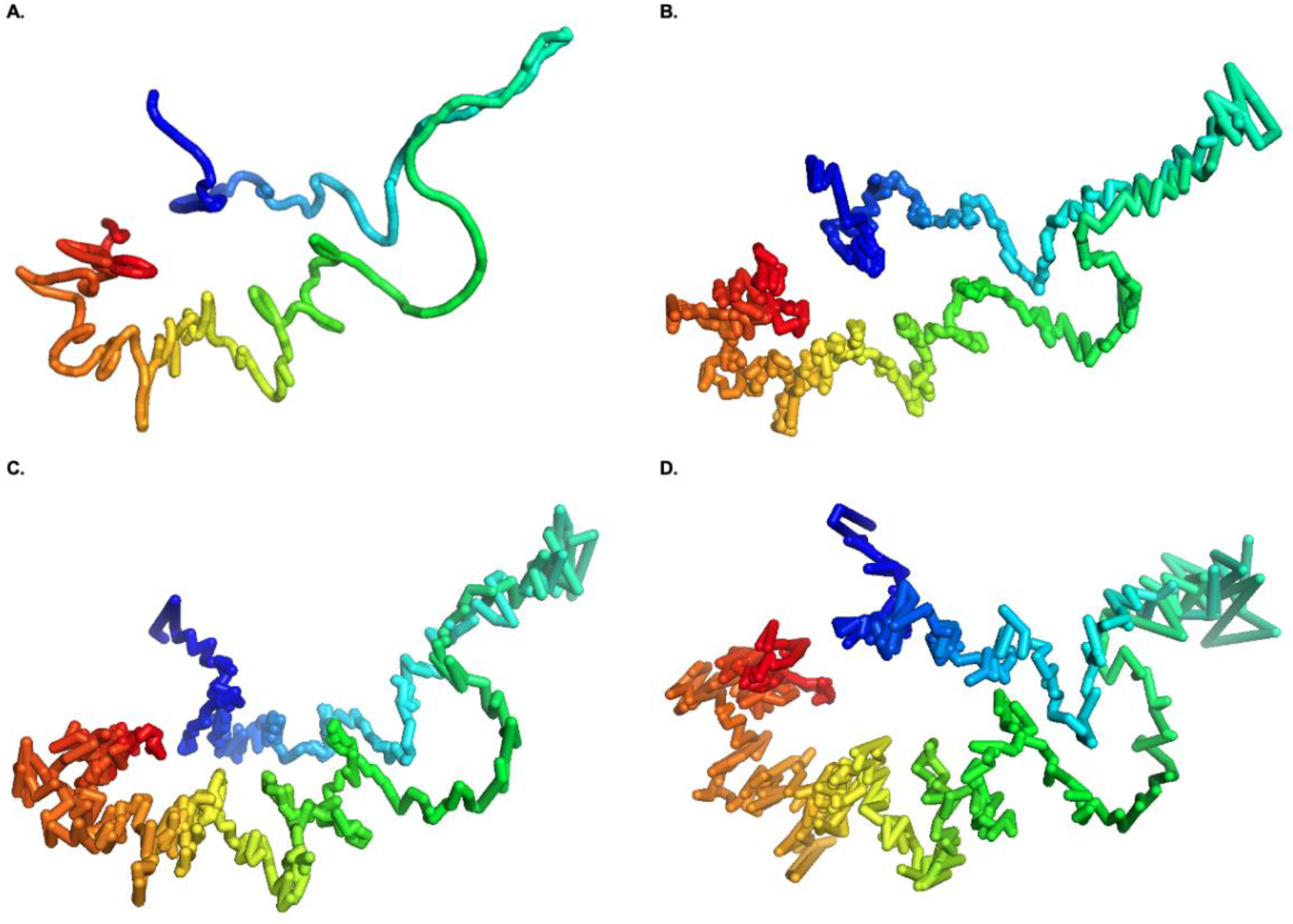
Structures generated by ParticleChromo3D at different Noise levels. Here, we show the structure generated by ParticleChromo3D at Noise level = (A) 0, that is no Noise, (B) 7% (70), (C) 20% (200), and (D) 40% (400)

### Assessment on Real Hi-C data

For evaluation on the real Hi-C data, we used the GM12878 B-lymphoblastoid cells line by Rao et al., 2014 [55]. The normalized 1MB and 500KB resolution interaction frequency matrices GM12878 cell line datasets were downloaded from the GSDB repository under the GSDB ID OO7429SF [56]. The datasets were normalized using the Knight-Ruiz normalization technique [15]. The performance of ParticleChromo3D was determined by computing the SCC value between the distance matrix of the normalized frequency input matrix and the Euclidean distance calculated from the predicted 3D structures. Fig 14 shows the assessment of ParticleChromo3D on the GM12878 cell line dataset. The reconstructed structure by ParticleChromo3D is compared against the input IF expected distance using the PCC, SCC, and RMSD metrics for the 1MB and 500KB resolution Hi-C data. When ParticleChromo3D performance is evaluated using both 1MB and 500KB resolution HiC data of the GM12878 cell, we observed some consistency in the algorithm’s performance for both datasets. Chromosome 18 had the lowest SCC value of 0.932 and 0.916 at 1MB and 500KB resolutions, respectively, while chromosome 5 had the highest SCC value of 0.975 and 0.966 at 1MB and 500KB resolutions, respectively.

**Fig 14.**
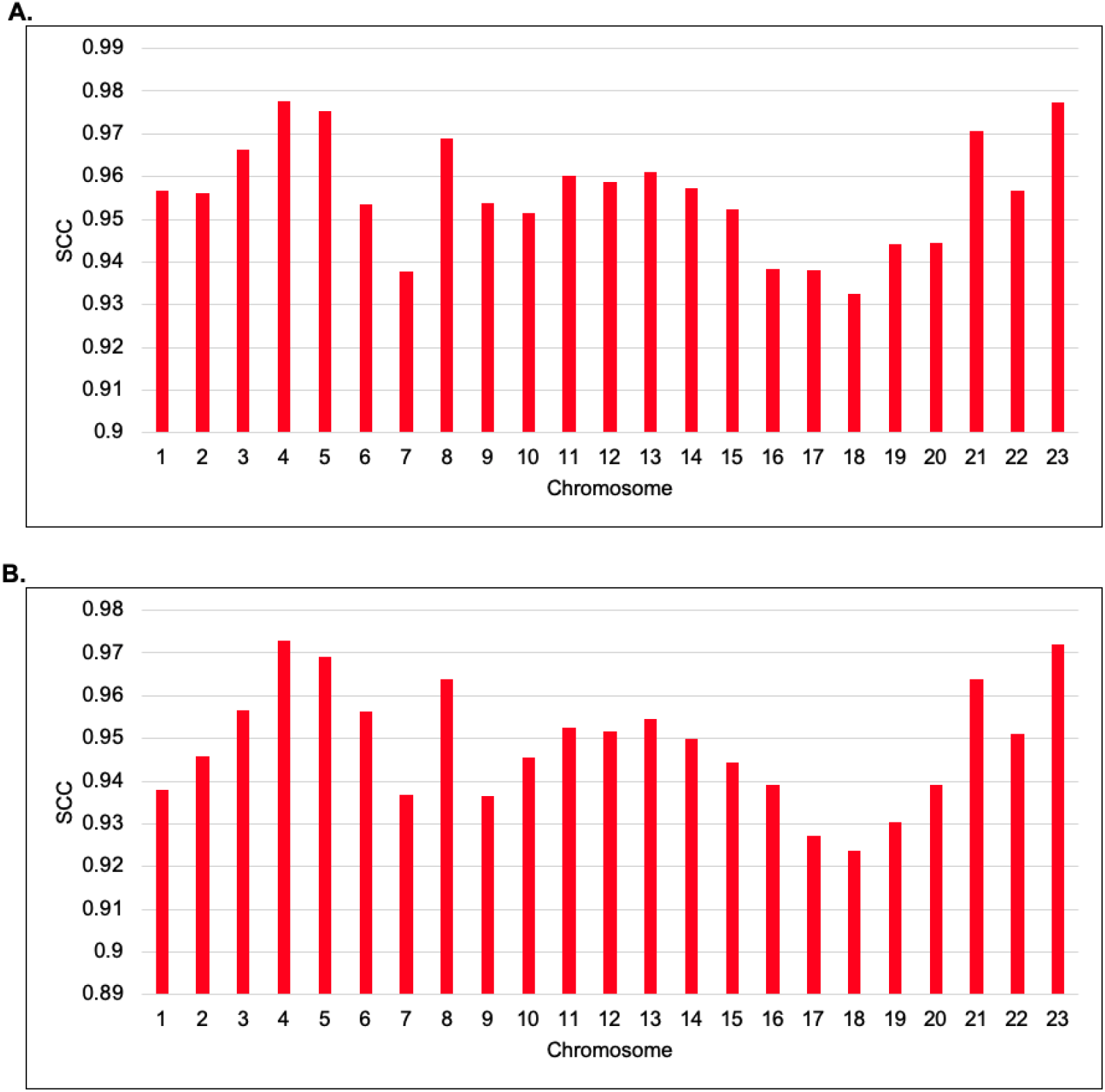
Performance evaluation of ParticleChromo3D using SCC values for 1MB and 500KB resolution GM12878 cell Hi-C data. (A) A plot of ParticleChromo3D SCC performance on 1MB GM12878 cell Hi-C data chromosome 1 to 23 (B) A plot of ParticleChromo3D SCC performance on 500KB GM12878 cell Hi-C data for chromosome 1 to 23.

### Model Consistency

Next, we assessed the consistency of our generated structures. We created 30 structures for the chromosomes and then evaluated the structure’s similarity using the SCC, PCC, RMSE, and TM-Score (Fig 15). We assessed the consistency for both the 1MB and 500KB resolution Hi-C data of the GM12878 cell. As illustrated for the TM-score, a score of 0.17 indicates pure randomness, and a score above 0.5 indicates the two structures have mostly the same folds. Hence the higher, the better. Our results show from the selected chromosomes that the structures generated by ParticleChromo3D are highly consistent for both the 1MB (Fig 15) and 500KB (Fig 16) datasets. As shown in Fig 15 for the 1MB Hi-C datasets, the average SCC and PCC values recorded between the models for the selected chromosomes is >=0.985 and >=0.988, respectively, indicating that chromosomal models generated by ParticleChromo3D are highly similar. It also indicates that it finds an absolute 3D model solution on each run of the algorithm (Fig 15C and Fig 15D). Similarly, as shown in Fig 16, for the 500KB Hi-C datasets, the average SCC and PCC values recorded between the models for the selected chromosomes is >= 0.992.

**Fig 15.**
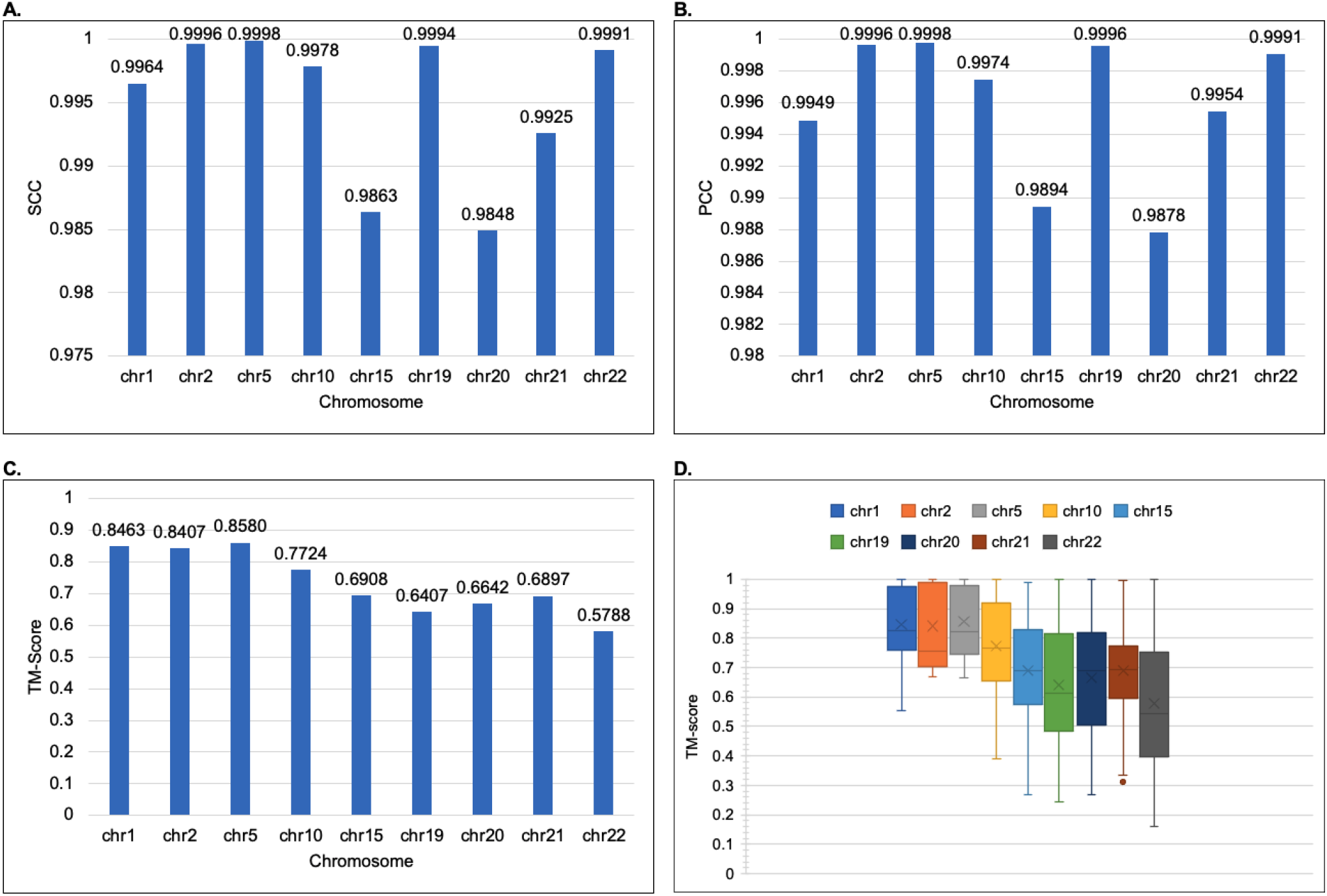
The model consistency check for 1MB resolution structures generated by ParticleChromo3D using different evaluation metrics. (A) The average SCC Between 30 Structures per chromosome at 1MB Resolution for the GM12878 datasets. (B) The average PCC Between 30 Structures per chromosome at 1MB Resolution for the GM12878 datasets.(C) The average TM-Score Between 30 Structure per chromosome at 1MB Resolution for the GM12878 datasets. (D) The boxplot shows the distribution of the 30 structure’s TM-score by chromosome for the GM12878 datasets. The Y-axis denotes the SCC and PCC metric score in the range [-1,1], and TM-Score in the range [-0,1]. The X-axis denotes the chromosome. A higher SCC, PCC, and TM-Score value is better.

**Fig 16.**
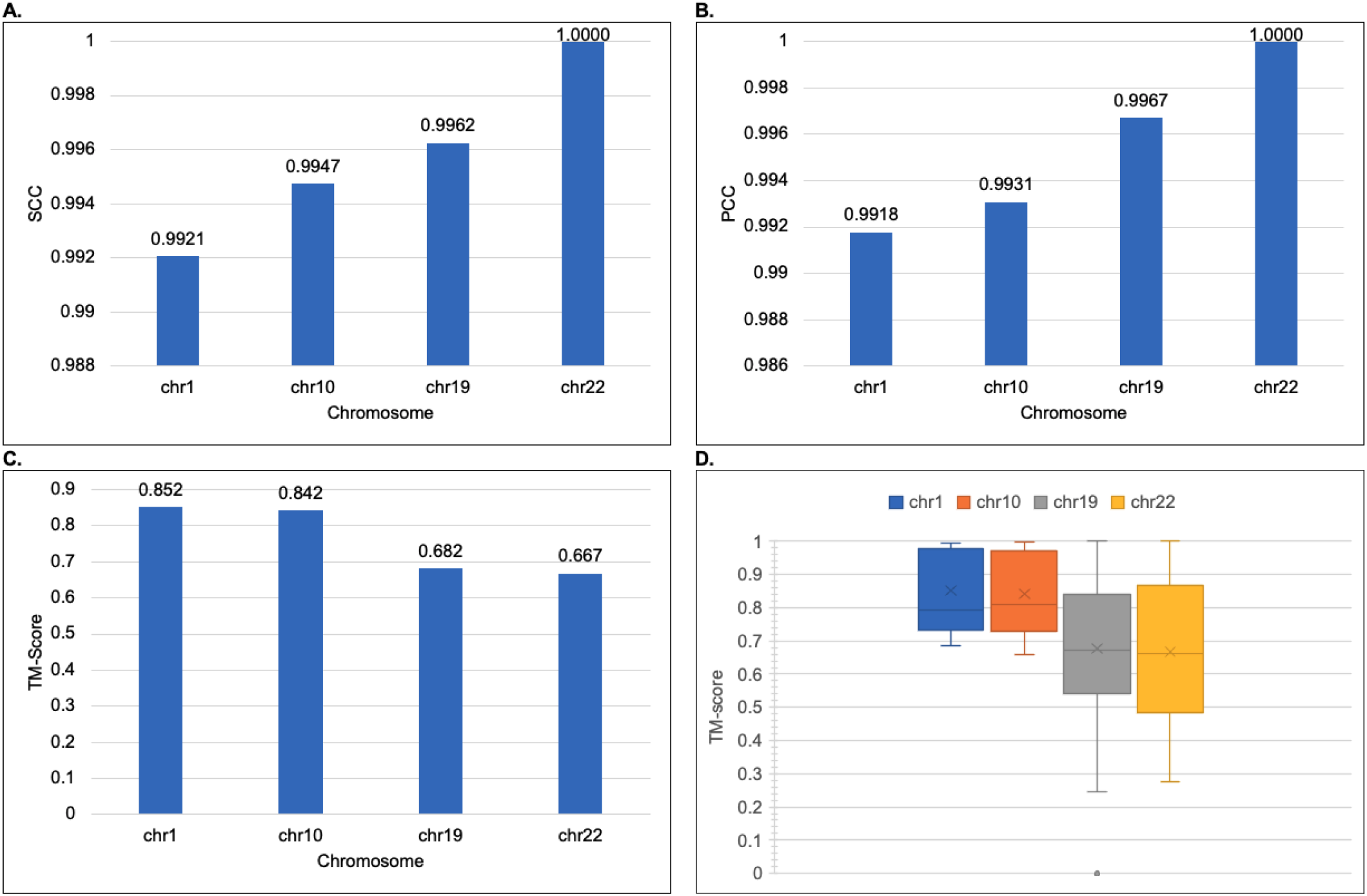
The model consistency check for 500KB resolution structures generated by ParticleChromo3D using different evaluation metrics. (A) The average SCC Between 30 Structures per chromosome at 500KB Resolution for the GM12878 datasets. (B) The average PCC Between 30 Structures per chromosome at 500KB Resolution for the GM12878 datasets. (C) The average TM-Score Between 30 Structure per chromosome at 500KB Resolution for the GM12878 datasets. (D) The boxplot shows the distribution of the 30 structure’s TM-score by chromosome for the GM12878 datasets. The Y-axis denotes the SCC and PCC metric score in the range [-1,1], and TM-Score in the range [-0,1]. The X-axis denotes the chromosome. A higher SCC, PCC, and TM-Score value is better.

### Comparison with existing chromosome 3D structure reconstruction methods

Here, we compared the performance of ParticleChromo3D side by side with nine existing high-performing chromosome 3D structure reconstruction algorithms on the GM12878 data set at both the 1MB and 500KB resolutions. The reconstruction algorithms are ChromSDE [26], Chromosome3D [28], 3DMax [29], ShRec3D [30], LorDG [31], GEM [72], HSA [33], MOGEN [35] and PASTIS [41] (Fig 17). According to the SCC value reported, we observed that ParticleChromo3D outperformed most of the existing methods in many chromosomes evaluated at 1MB and 500KB resolution. At a minimum, ParticleChromo3D secured the top-two best overall performance position among the ten algorithms compared. ParticleChromo3D achieving these results against these methods and algorithms shows the robustness and suitability of the PSO algorithm to be used to solve the 3D chromosome and genome structure reconstruction problem.

**Fig 17.**
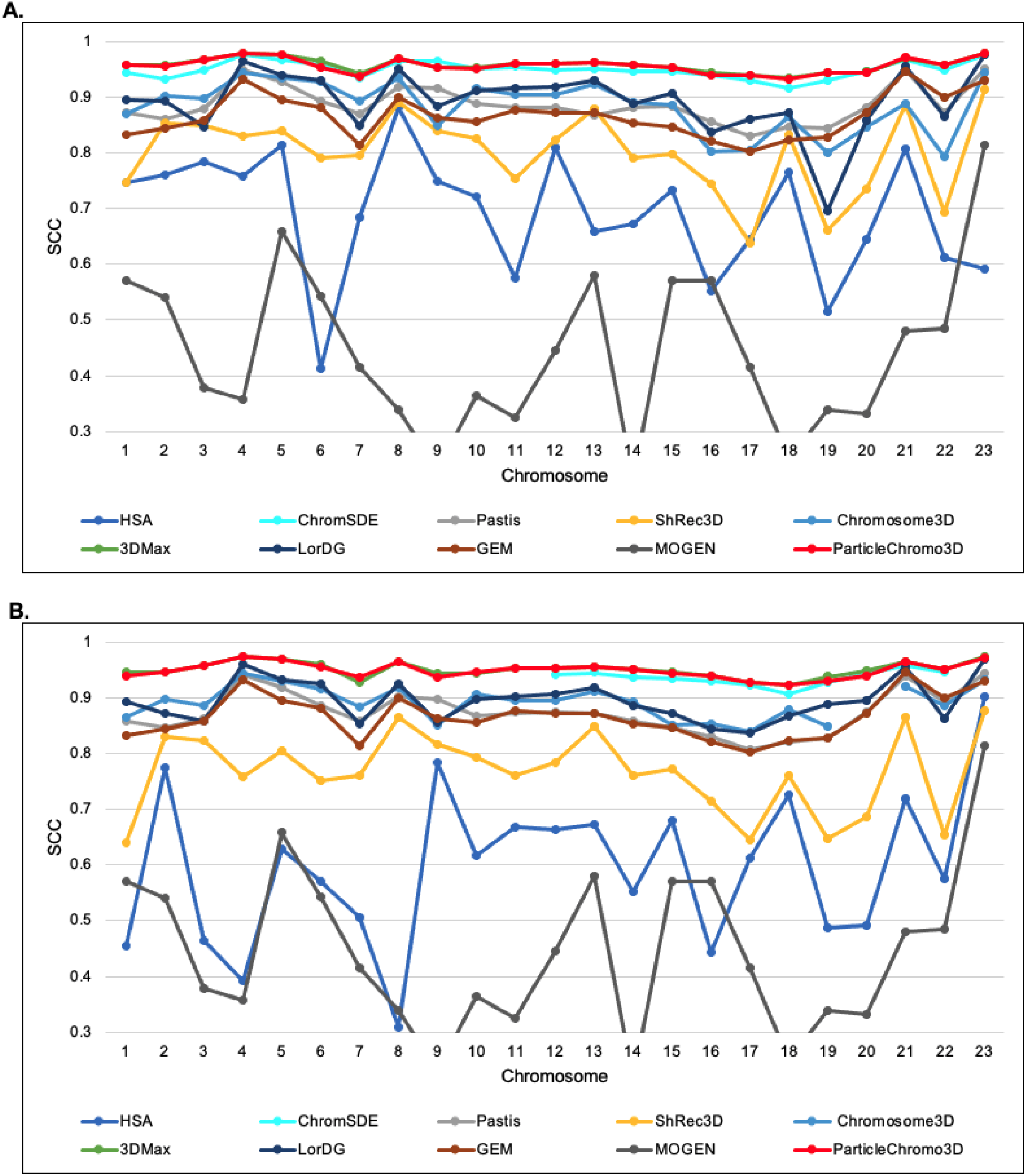
A comparison of the accuracy of nine existing methods and ParticleChromo3D for 3D structure reconstruction on the 1MB and 500KB real Hi-C dataset. (A) An SCC Comparison of 3D structure reconstruction methods on the GM12878 Hi-C dataset at 1Mb resolution for chromosomes 1 to 23. (B) An SCC Comparison of 3D structure reconstruction methods on the GM12878 Hi-C dataset at 500KB resolution for chromosomes 1 to 23. The Y-axis denotes the SCC metric score in the range [-1,1], and X-axis denotes the chromosome. A higher SCC value is better.

## Discussion

We discussed the Swarm Size value’s relevance in the Parameters Estimation section. We showed on the synthetic dataset that a Swarm Size (SS) value of 5 did not produce satisfactory performance. However, it was the fastest considering the other swarm sizes. At SS = 10, the performance was significantly improved than at SS = 5, but with an increase in computation time as a consequence. SS values 15 and 20 similarly achieved better performance, but the cost of this performance improvement similarly is an increase in the program running time. However, we settled for a SS = 15 because it achieved one of the best performances, and the computational cost can be considered manageable. To investigate the implication of our choice, we carried out two tests discussed below:

### ParticleChromo3D performance on different Swarm Size values

First, we evaluated the performance of the ParticleChromo3D algorithm on the GM12878 data set on both the 1MB and 500KB resolutions at Swarm Sizes 5, 10, and 15 to ensure that the performance at SS = 15 that we observed on the synthetic dataset is carried over to the real dataset (Fig 18). The 1MB and 500KB dataset result shows that SS = 15 achieved the best SCC value mostly across the chromosomes (Fig 18). However, we observed that the result generated at SS = 10 were also competitive and achieved an equal performance a few times with SS = 15. This shows us that choosing the SS = 10 does not necessarily reduce the performance of our ParticleChromo3D. There is an additional gain of saving on computational time if this value is used.

**Fig 18.**
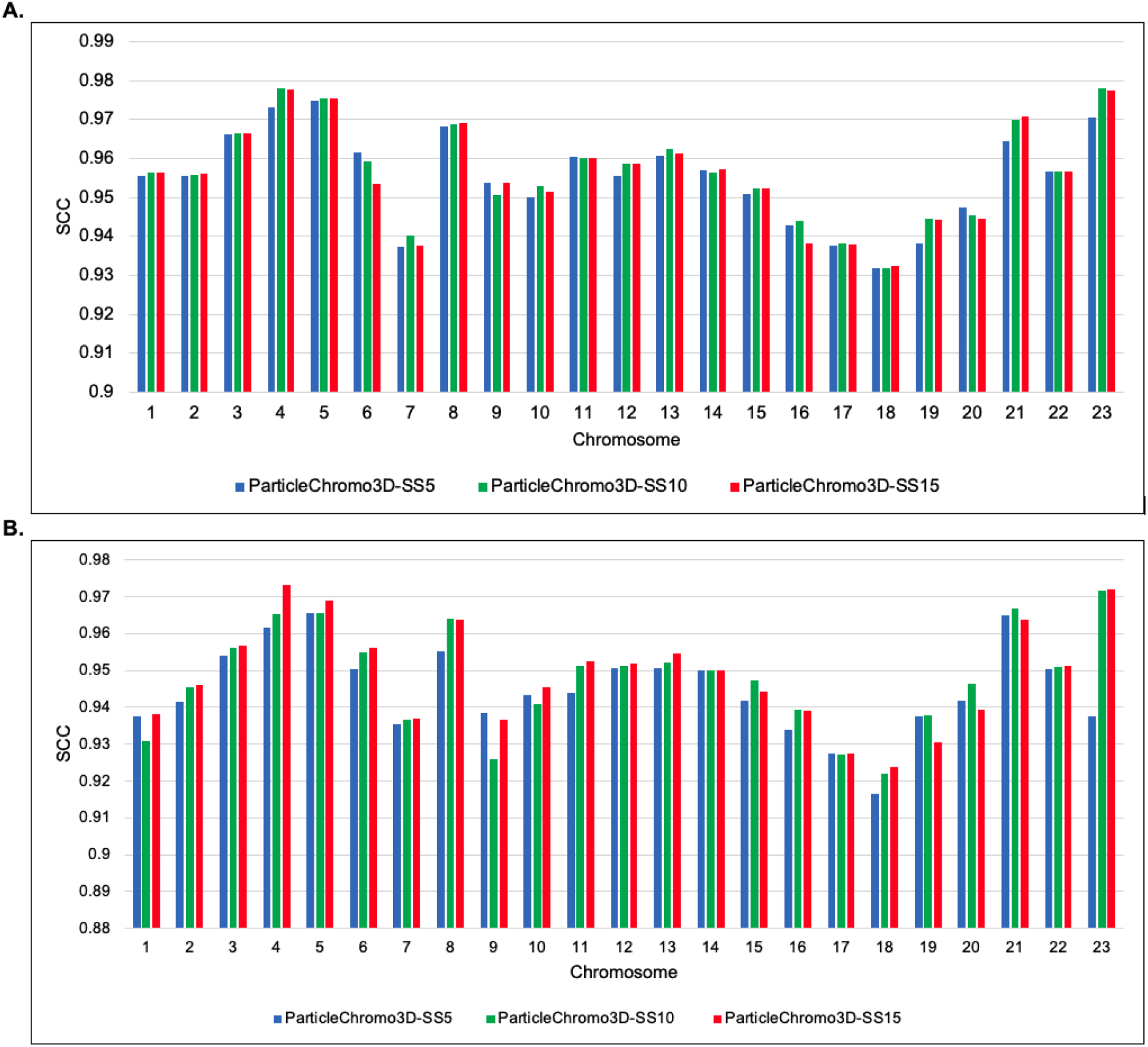
ParticleChromo3D SCC performance on Swarm Size values 5,10 and 15 for 1MB and 500KB GM12878 cell Hi-C data. (A) Comparing the performance by ParticleChromo3D on the 1MB GM12878 cell Hi-C data at Swarm Size values 5, 10, and 15. (B) Comparing the performance by ParticleChromo3D on the 500KB GM12878 cell Hi-C data at Swarm Size values 5, 10, and 15. The Y-axis denotes the SCC metric score in the range [-1,1], and X-axis denotes the chromosome. A higher SCC value is better.

### Computational Time

Second, we evaluated the time it took our algorithm to perform the 3D reconstruction for select chromosomes of the 1MB and 500KB GM12878 cell Hi-C data set. The modeling of the structures generated by ParticleChromo3D for the synthetic and real dataset was done on an AMD Ryzen 7 3800x 8-Core Processor, 3.89GHZ with installed RAM 31.9GB.

ParticleChromo3D is programmed to multithread. It utilizes each core present on the user’s computer to run a specific task, speeding up the modeling process and significantly reducing computational time. Accordingly, the more the number of processors a user has, the faster ParticleChromo3D will generate an output 3D structures. As mentioned earlier in the Parameter Estimation section, one of the default settings for ParticleChromo3D is to automatically determine the best conversion factor that fits the data in the range [0.1, 1.5]. Even though this is one of our ParticleChromo3D’s strengths, this process has the consequence of increasing the algorithm’s computational time. Based on the real Hi-C dataset analysis, our result shows that the Swarm Size 10 consistently has a lower computational time than the SS = 15 as speculated for the 500KB and 1MB Hi-C datasets (Fig 19). These results highlight an additional strength of ParticleChromo3D that it can achieve a competitive result in a lower time (Fig 19) without trading it off with performance (Fig 18). It is worth noting that we recommend that users can set the Swarm Size to the preferred value depending on the objective. In this manuscript, we favored the algorithm achieving a high accuracy over speed. We made up for this by making our algorithm multi-threaded, reducing the running time significantly.

**Fig 19.**
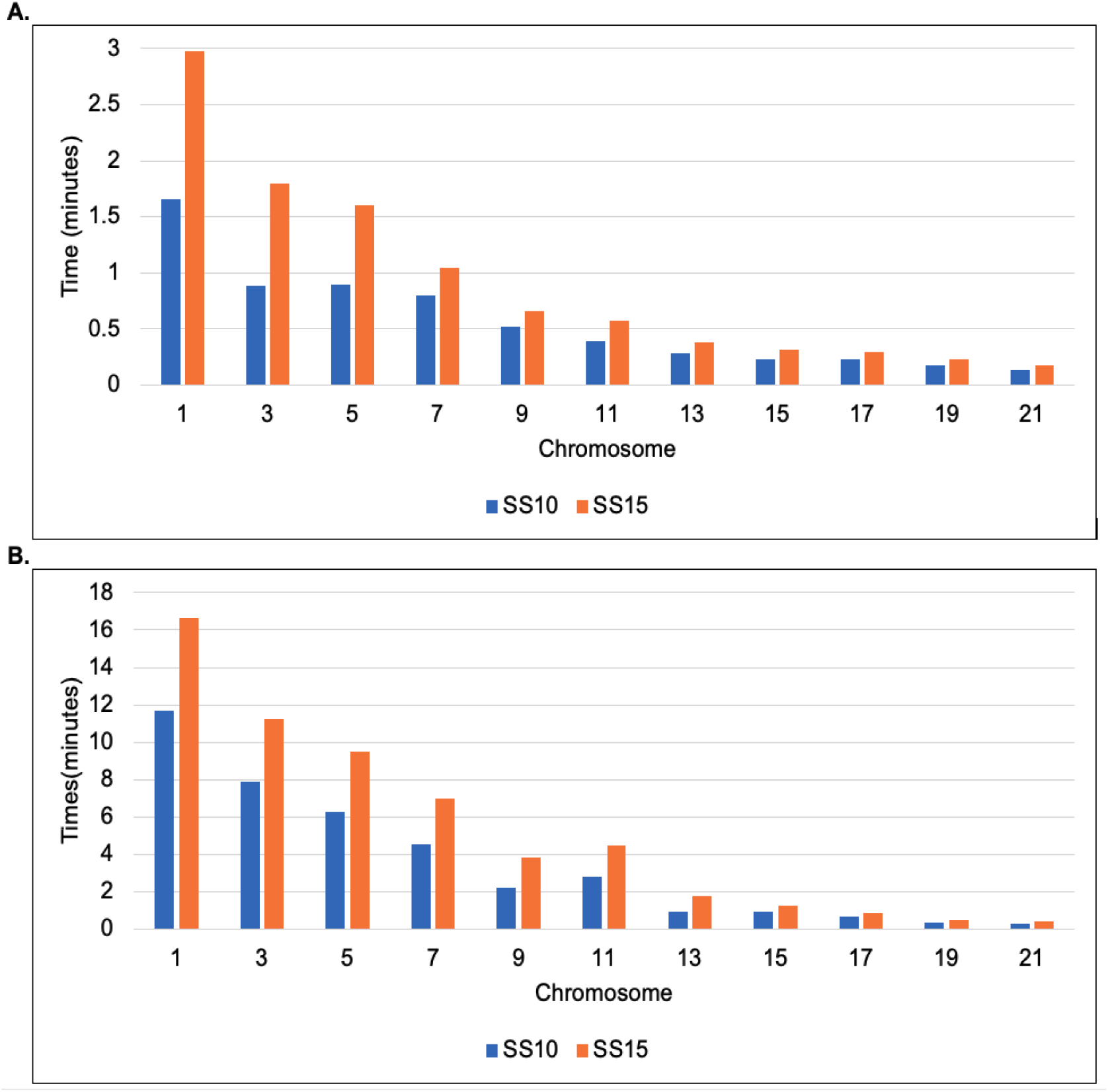
ParticleChromo3D Computational Time at Swarm Size (SS) 10 and 15 for 1MB and 500KB GM12878 cell Hi-C data. (A) Comparing the running time for ParticleChromo3D for select chromosomes for 1MB GM12878 cell Hi-C data (B)A comparison of the running time for ParticleChromo3D for select chromosomes for 500KB GM12878 cell Hi-C data. The Y-axis denotes the running time for ParticleChromo3D in minutes, and X-axis denotes the chromosome.

### Availability of data and Materials

The models generated, all the datasets used for all analysis performed, and the source code for ParticleChromo3D are available at https://github.com/OluwadareLab/ParticleChromo3D.

## Conclusions

We developed a new algorithm for 3D genome reconstruction called ParticleChromo3D. ParticleChromo3D uses the Particle Swarm Optimization algorithm as the foundation of its solution approach for 3D chromosome reconstruction from Hi-C data. The results of ParticleChromo3D on simulated data show that with the best-fine-tuned parameters, it can achieve high accuracy in the presence of noise. We compared ParticleChromo3D accuracy with nine (9) existing high-performing methods or algorithms for chromosome 3D structure reconstruction on the real dataset. The results show that ParticleChromo3D is effective and a high performer by achieving more accurate results over the other methods in many chromosomes; and securing the top-two best overall position in our comparative analysis with other algorithms. Our experiments also show that ParticleChromo3D can also achieve a faster computational run time without losing accuracy significantly. ParticleChromo3D’s parameters have been optimized to achieve the best result for any input Hi-C by searching for the best conversion factor (*α*) and using the optimal PSO hyperparameters for any given input automatically. This algorithm was implemented in python and can be run as an executable or as a Jupyter Notebook found at https://github.com/OluwadareLab/ParticleChromo3D.

## Acknowledgments

Not applicable.

## Supporting information

**S1 Fig. A Plot of local confidence values 0.3 and 0.5 versus global confidence.**

(A) SCC by Global Confidence at Local Confidence 0.3 (B) PCC by Global Confidence at Local Confidence 0.3. (C) SCC by Global Confidence at Local Confidence 0.5 (D) PCC by Global Confidence at Local Confidence 0.5. Each of the plots shows the SCC and PCC results obtained by comparing the ParticleChromo3D algorithm’s output structure with the simulated dataset’s true structure for local confidence values 0.3 to 0.5 and global confidence values 0.1 to 2.8. The Y-axis denotes the SCC or PCC scores, respectively, as a label in the title, in the range [-1,1], the X-axis denotes the global confidence values, and the colored plot denotes the local confidence values. A higher SCC and PCC value is better.

**S2 Fig. A Plot of local confidence values 0.7 and 0.9 versus global confidence.**

(A) SCC by Global Confidence at Local Confidence 0.7. (B) PCC by Global Confidence at Local Confidence 0.7. (C) SCC by Global Confidence at Local Confidence 0.9. (D) PCC by Global Confidence at Local Confidence 0.9. Each of the plots shows the SCC and PCC results obtained by comparing the ParticleChromo3D algorithm’s output structure with the simulated dataset’s true structure for local confidence values 0.7 to 0.9 and global confidence values 0.1 to 2.8. The Y-axis denotes the SCC or PCC scores, respectively, as a label in the title, in the range [-1,1], the X-axis denotes the global confidence values, and the colored plot denotes the local confidence values. A higher SCC and PCC value is better.

