## Supplementary figures and images for "ParticleChromo3D: A Particle Swarm Optimization Algorithm for Chromosome and Genome 3D Structure Prediction from Hi-C Data"

### S1 Fig

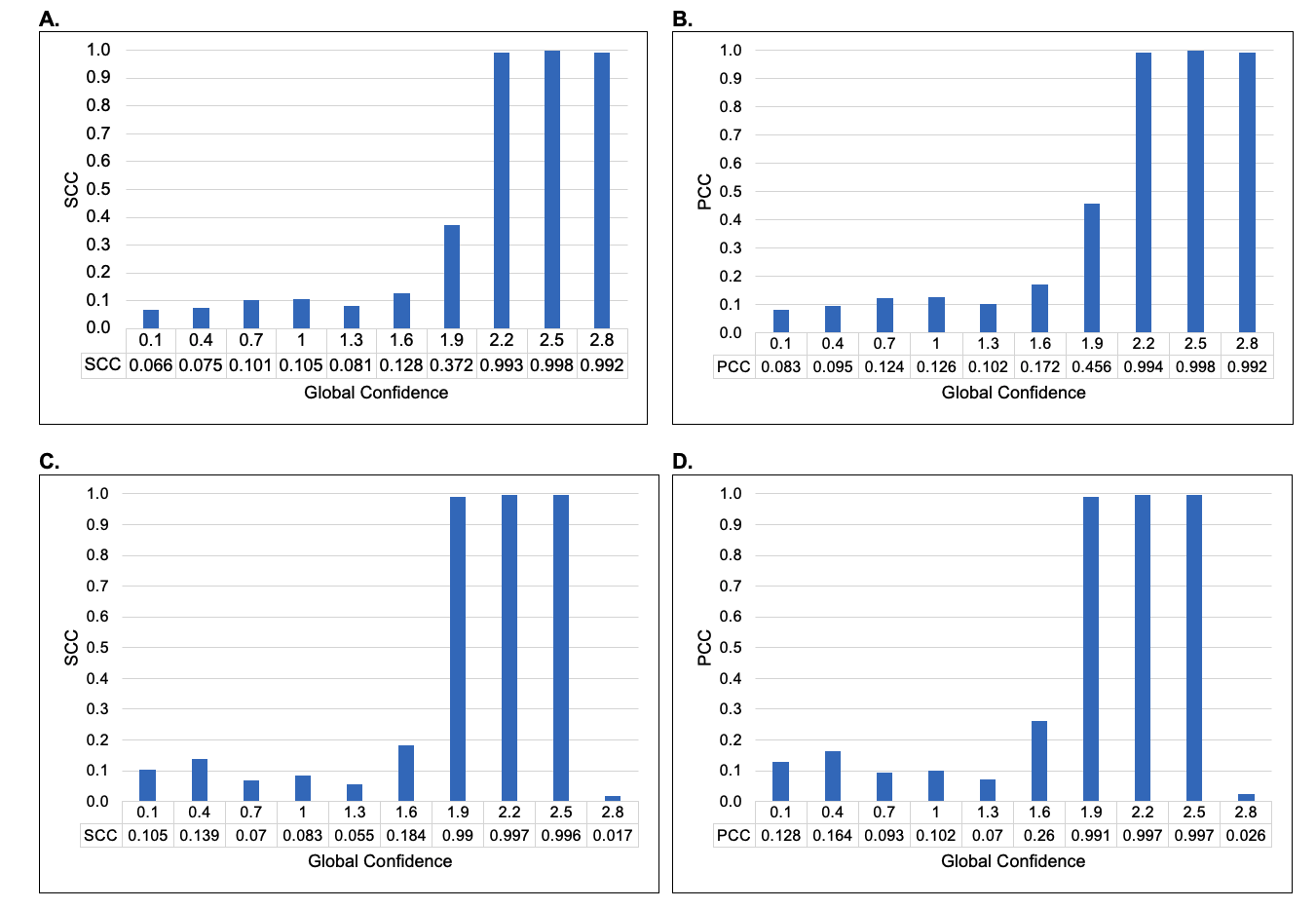

### S2 Fig

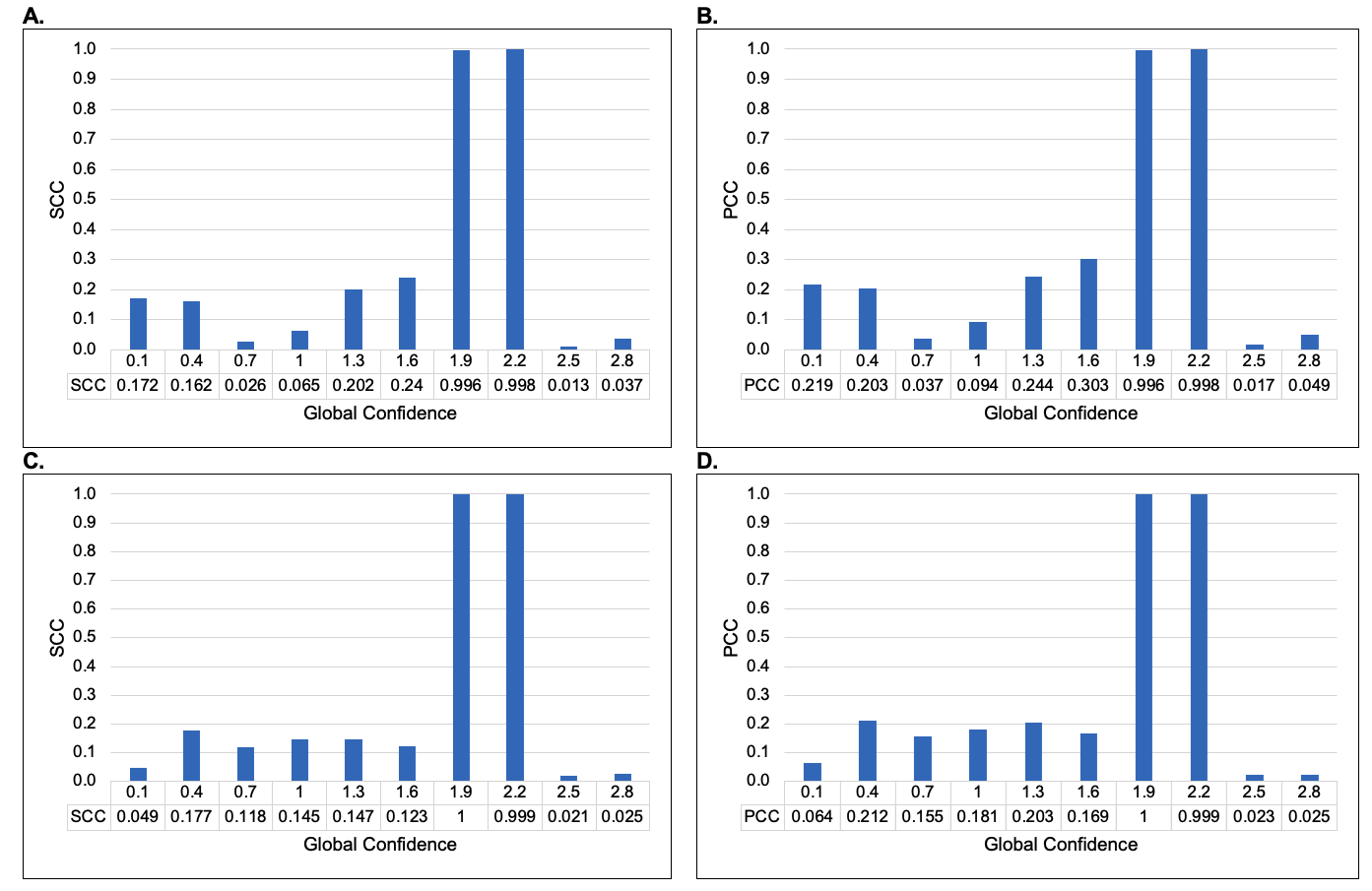
